# Sterol flow between the plasma membrane and the endosome is regulated by the LAM family protein Ltc1

**DOI:** 10.1101/720383

**Authors:** Magdalena Marek, Vincent Vincenzetti, Sophie G. Martin

## Abstract

Sterols are crucial components of biological membranes that help maintain membrane integrity and regulate various processes such as endocytosis, protein oligomerization and signaling. Although synthetized in the ER, sterols are at highest concentrations at the plasma membrane (PM) in all eukaryotic organisms. Here, by applying a genetically encoded sterol biosensor (D4H), we visualize a sterol flow between PM and endosomes in the fission yeast *Schizosaccharomyces pombe*. While D4H is detected at the PM during steady-state growth, inhibition of Arp2/3-dependent F-actin assembly unexpectedly promotes the reversible re-localization of the probe to internal sterol rich compartments (STRIC), as shown by correlative light-electron microscopy. Time-lapse imaging identifies STRIC as a late secretory, endosomal compartment labelled by the synaptobrevin Syb1. Retrograde sterol internalization to STRIC is independent of endocytosis or an intact Golgi. Instead, it depends on Ltc1, a LAM/StARkin-family protein that localizes to ER-PM contact sites. In *ltc1Δ*, sterols over-accumulate at the PM, which forms extended ER-interacting invaginations, indicating that sterol transfer by Ltc1 contributes to PM size homeostasis. Anterograde sterol movement from STRIC is independent of canonical vesicular trafficking components but requires Arp2/3 activity, suggesting a novel physiological role for this complex. Thus, transfer routes orthogonal to vesicular trafficking govern the retrograde and anterograde flow of sterols in the cell.

## Introduction

Sterols are critical components of biological membranes with fundamental roles in cellular physiology. They confer essential biophysical properties to bio-membranes, regulate vesicular trafficking and signal transduction. Moreover, they serve as precursors of hormones, bile acids, vitamin D and energy storage molecules. While sterols can be either synthesized in cells or taken up from the environment, dysfunctions in their trafficking or metabolism cause severe pathologies, such as Niemann-Pick disease type C, hypercholesterolemia or sitosterolemia. In fungi, the sterol molecule ergosterol is essential for viability and serves as a major target of antifungal drugs.

Although sterols are synthesized in the endoplasmic reticulum (ER), their concentration within the ER does not exceed 0.5-1 mol% of total lipids (Luo et al., 2017). Instead, it increases along the secretory pathway reaching about 40 mol% at the plasma membrane (PM) (Maxfield and van Meer, 2010). Within the PM, sterols cluster with other lipids, especially sphingolipids, forming membrane subdomains that range from the nanoscale to several μm (Alvarez et al., 2007; Makushok et al., 2016; Sezgin et al., 2017).

How sterol domains are formed and maintained, and how sterols are transported between different organelles has been the subject of intense investigation. Previous work in mammalian and yeast cells showed that the transport of newly synthetized cholesterol from the ER to the PM is fast, with a half-time of a few minutes (Baumann et al., 2005; Georgiev et al., 2011), and does not occur through the canonical secretory pathway. Indeed, collapse of the Golgi apparatus by Brefeldin A treatment in mammalian cells or secretory pathway block with ts mutants in yeast has no or little effect on sterol transport rates (Baumann et al., 2005; Heino et al., 2000; Urbani and Simoni, 1990). Instead, sterol molecules are thought to be carried between membranes by Lipid Transfer Proteins (LTPs) (Wong et al., 2017).

We can distinguish two major classes of proteins implicated in sterol transport (Luo et al., 2019). The first is the ORP family (<u>O</u>xy<u>s</u>terol <u>b</u>inding <u>p</u>rotein (OSBP)-Related Protein) known in *S. cerevisiae* as OSH (<u>O</u>xy<u>s</u>terol-<u>b</u>inding <u>p</u>rotein Homologue). Not all proteins in this family transport sterol, but they bind phosphoinositides as common ligand, leading to the hypothesis that some ORPs (e.g. mammalian OSBP and *S. cerevisiae* Osh4/Kes1) transport sterol between membranes by counter exchange with phosphoinositides (Antonny et al., 2018). Surprisingly, knockout of all seven OSH proteins in yeast did not significantly affect bidirectional sterol transport, suggesting that either OSH’s primary function is not bulk sterol transport, or that there exists a large degree of redundancy with other sterol transport pathways (Georgiev et al., 2011; Raychaudhuri et al., 2006).

The second class of sterol transport proteins is part of the large STARkin superfamily, which possess a StART (Steroidogenic Acute Regulatory Transfer) or StART-like lipid-binding domain (Wong and Levine, 2016). The StART domain, able to bind sterols, phospho- and/or sphingolipids, define the STARD family, which is absent from fungi and Archea. The StART-like domain, also known as VASt (VAD1 Analog of StART) (Khafif et al., 2014), binds cholesterol and ergosterol, and promotes the transfer of sterol molecules between membranes (Gatta et al., 2018; Horenkamp et al., 2018; Jentsch et al., 2018; Tong et al., 2018). In *S. cerevisiae*, this protein family, named LAM (Lipid transfer proteins Anchored at Membrane contact sites), consists of proteins typically anchored in the ER membrane through a transmembrane domain and enriched at organelle contact sites (Elbaz-Alon et al., 2015; Gatta et al., 2015; Murley et al., 2015). Lam1-4 localize to ER-PM contact sites, with Lam2/Ysp2 involved in retrograde sterol transport (Gatta et al., 2015); Lam6/Ltc1 localizes to contact sites between ER, mitochondria and vacuoles (Elbaz-Alon et al., 2015; Murley et al., 2015); Lam5/Ltc2 may localize to contact sites between the ER and the late Golgi (Weill et al., 2018). The mammalian homologues, variously called GramD1 or Aster, also localize to ER-PM contact sites and contribute to retrograde transport of exogenously provided cholesterol (Besprozvannaya et al., 2018; Sandhu et al., 2018). Intriguingly, while most LTPs localize to dedicated organelle contact sites, *S. cerevisiae* mutants lacking all six known ER-PM tethering proteins show normal anterograde, and only partially diminished retrograde sterol transport (Quon et al., 2018).

Yeast cells, such as *S. cerevisiae* and *S. pombe*, have simple endomembrane system. For instance, the trans-Golgi network (TGN) in *S. cerevisiae* also serves as early and late endosomal compartment (Day et al., 2018). Both organisms harbour a single, well-characterized, endocytic pathway that uses Arp2/3-mediated actin assembly to generate forces for internalization and retrieval of proteins from the PM to the TGN (Goode et al., 2015; Lacy et al., 2018). Vesicular trafficking from the TGN to the PM relies on transport along actin cables, assembled by the formin For3 in fission yeast, and tethering at the PM by the exocyst complex (Bendezu and Martin, 2011; Feierbach et al., 2004; TerBush et al., 1996; Wang et al., 2002). Other complexes such as exomer or clathrin adaptors contribute to sorting from the TGN (Hoya et al., 2017; Wang et al., 2006). Our extended knowledge of protein trafficking contrasts sharply with that of sterol transport, which remains mostly unknown.

A major difficulty in studying intracellular sterol transport lies in its visualization. The fluorescent antibiotic filipin, a widely used polyene macrolide-based sterol dye, is not suitable for prolonged live imaging due to toxicity (Alvarez et al., 2007). Other approaches rely on application of fluorescently labelled sterol analogues, which however carry bulky chemical groups that can alter sterol behavior (Sezgin et al., 2016). Moreover, yeast cells only take them up in specific mutant backgrounds or growth conditions (Jacquier and Schneiter, 2012; Marek et al., 2014). Domain 4 (D4) of the bacterial toxin perfringolysin O (PFO), which binds membranes containing ≥30 mol% sterols and can be expressed in cells with no obvious toxicity, offers an interesting alternative (Shimada et al., 2002). Part of the reason for the high detection threshold is that PFO detects free sterol in the membrane, but not sterol in complex with sphingolipids (Das et al., 2014). An improved allele, called D4H, capable of binding membranes with 20 mol% sterols (Johnson et al., 2012; Maekawa and Fairn, 2015) represents a promising genetically-encoded biosensor for tracking sterols *in vivo* (Maekawa et al., 2016).

Here, we applied the D4H probe to visualize sterol-rich membranes in the fission yeast *Schizosaccharomyces pombe* and discover a route of massive sterol internalization from the PM, triggered by depolymerization of the actin cytoskeleton. We show that sterols undergo retrograde trafficking from the PM to endosomes, in a manner strictly dependent on a single, previously uncharacterized StARTkin-protein Ltc1, and return to the PM by means independent of vesicular trafficking.

## Results

### D4H as a bio-probe for intracellular sterol visualization

To visualize the intracellular distribution of sterols in live fission yeast cells, we expressed the mCherry-D4H sensor integrated as single copy under control of the constitutive actin promoter (*p*^*act1*^). Cells expressing this construct did not exhibit growth problems or morphological defects (Fig S1A and S1B). The biosensor localized to the cell periphery throughout the cell cycle, with occasional occurrence of one or a few internal dots (Fig 1A, Movie S1). The un-mutagenized mCherry-D4 probe also localized to the cell cortex, but exhibited 3-fold lower cortex-to-cytosol signal ratio, consistent with its lower sterol affinity (Fig 1B). Treatment of cells with the squalene epoxidase inhibitor terbinafine, which blocks an early step in sterol synthesis, led to D4H delocalization to the cytosol confirming the probe’s sterol-binding specificity (Fig 1C). Note that terbinafine treatment was cytostatic and did not lead to substantial cell death (Fig S1C-D-E). Restart of sterol biosynthesis upon terbinafine removal led to return of D4H to the cortex (Fig 1D). We conclude that the D4H sensor reports on the distribution of sterol-rich membranes in yeast cells.

**Figure 1.**
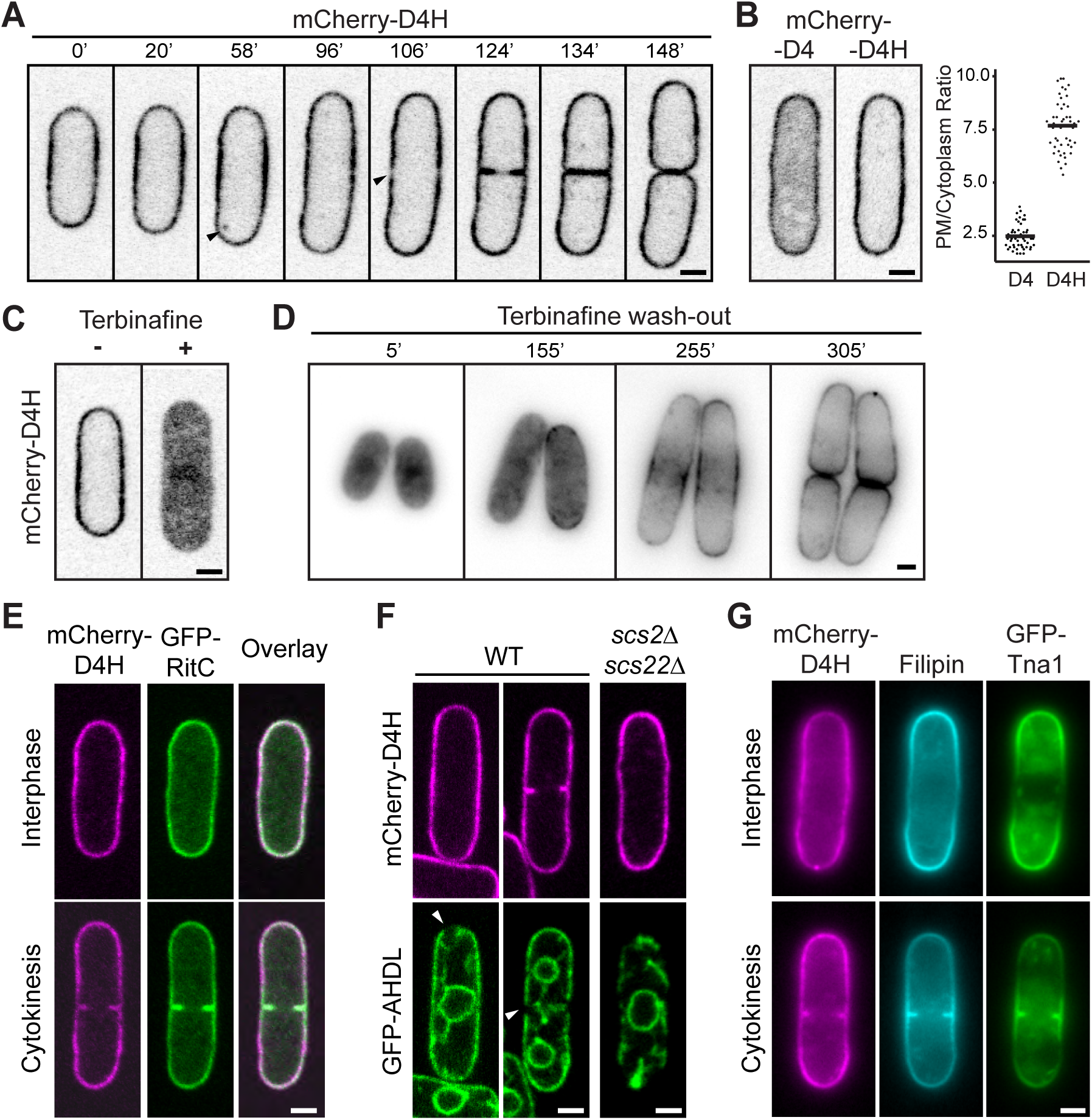
The D4H sterol bio-sensor labels the plasma membrane in fission yeast cells. **A.** mCherry-D4H distribution during interphase growth in *S. pombe* cells. The arrowheads point to an internal dot at 58’ and clearance of D4H from the pre-divisional site at 106’. Time in minutes. **B.** Localization of D4H and D4 sterol biosensors in vegetative cells. The plot on the right shows the PM to cytoplasm fluorescence ratio (n > 50 cells). Horizontal bars indicate the mean. **C.** mCherry-D4H distribution in cells treated with the sterol biosynthesis inhibitor terbinafine for 16h. **D.** Recovery of mCherry-D4H PM localization upon wash-out of terbinafine after 16h treatment. **E.** mCherry-D4H colocalizes with the PM marker GFP-RitC. **F.** mCherry-D4H decorates the PM, not the ER marked with GFP-AHDL in WT and ER-PM attachment-deficient mutant (*scs2Δ scs22Δ*). Arrowheads point to zones of PM devoid of ER. **G)** Distinct distribution of mCherry-D4H and filipin staining in cells also co-expressing GFP-Tna1. Scale bars 2μm.

To probe whether the D4H signal was at the PM or in the cortical ER, we co-expressed mCherry-D4H with the PM marker GFP-RitC (Bendezu et al., 2015) and ER marker GFP-AHDL. GFP-RitC fully colocalized with mCherry-D4H at the cell periphery and at the division site (Fig 1E). Consistent with the close ER-PM apposition, GFP-AHDL and mCherry-D4H also showed extensive colocalization, but specific regions at cell tips and the division plane labelled by D4H were devoid of ER (Fig 1F). Moreover, in mutants lacking VAP-family ER-PM tethers (*scs2Δ scs22*Δ), in which the ER is largely detached from the PM (Zhang et al., 2012), mCherry-D4H remained at the cell periphery (Fig 1F). Thus, D4H localizes to the PM, in line with the known high concentration of sterols in this compartment.

Interestingly, D4H intensity varied along the PM, between homogeneous distribution to slight depletion at cell poles during polarized growth. Similarly, the signal cleared from the cell division site shortly before cell division (Fig 1A). This distribution is distinct from that of filipin, which is enriched at cell poles and division site (Fig 1G) (Wachtler et al., 2003). The difference in D4H and filipin distributions may arise from distinct sterol distribution or different sterol accessibility in the inner and outer PM leaflets. D4H localization was also distinct from that of GFP-Tna1, a multipass transmembrane protein proposed to mark sterol-rich membrane domains (Makushok et al., 2016).

### Arp2/3 inhibition results in intracellular sterol accumulation

We unexpectedly discovered that depolymerization of F-actin by Latrunculin A (LatA) treatment caused mCherry-D4H loss from the PM and intracellular accumulation as discrete dots (Fig 2A). Specific Arp2/3 inhibition with CK-666, which depolymerizes endocytic actin patches but not actin cables, had the same effect (Fig 2A, Movie S2) (Nolen et al., 2009). CK-666 led to a transient D4H enrichment at cell poles, appearance of a weak diffuse signal, formation of D4H-positive structures within 15 min, and complete clearance from the PM within 30 to 60 min (Fig 2B-C). Cells pre-treated with CK-666 for 1h became resistant to further treatment with amphotericin B (AmB), an ergosterol-binding antifungal that induces the formation of pores in sterol-rich membranes (Fig 2D), in agreement with a reduction in PM sterols. Moreover, the PM of these cells did not stain with filipin (Fig 2E). Thus Arp2/3 inhibition leads to strong reduction in PM sterol levels. The effects of CK-666 were fully reversible: 30 to 60 min after wash out, D4H decorated again the PM (Fig 2B lower panel, Movie S3). Inhibition of Arp2/3 function by using the *arp2-1* temperature-sensitive allele yielded similar signal internalization (Fig 2F). By contrast, D4H remained at the PM upon formin For3 deletion, though this mutant exhibited more internal dots than wildtype in untreated growth conditions (Fig 2G). Microtubule depolymerization had no effect on mCherry-D4H distribution (Fig S2). These results indicate a crucial role for F-actin assembly by Arp2/3 in sterol distribution.

**Figure 2.**
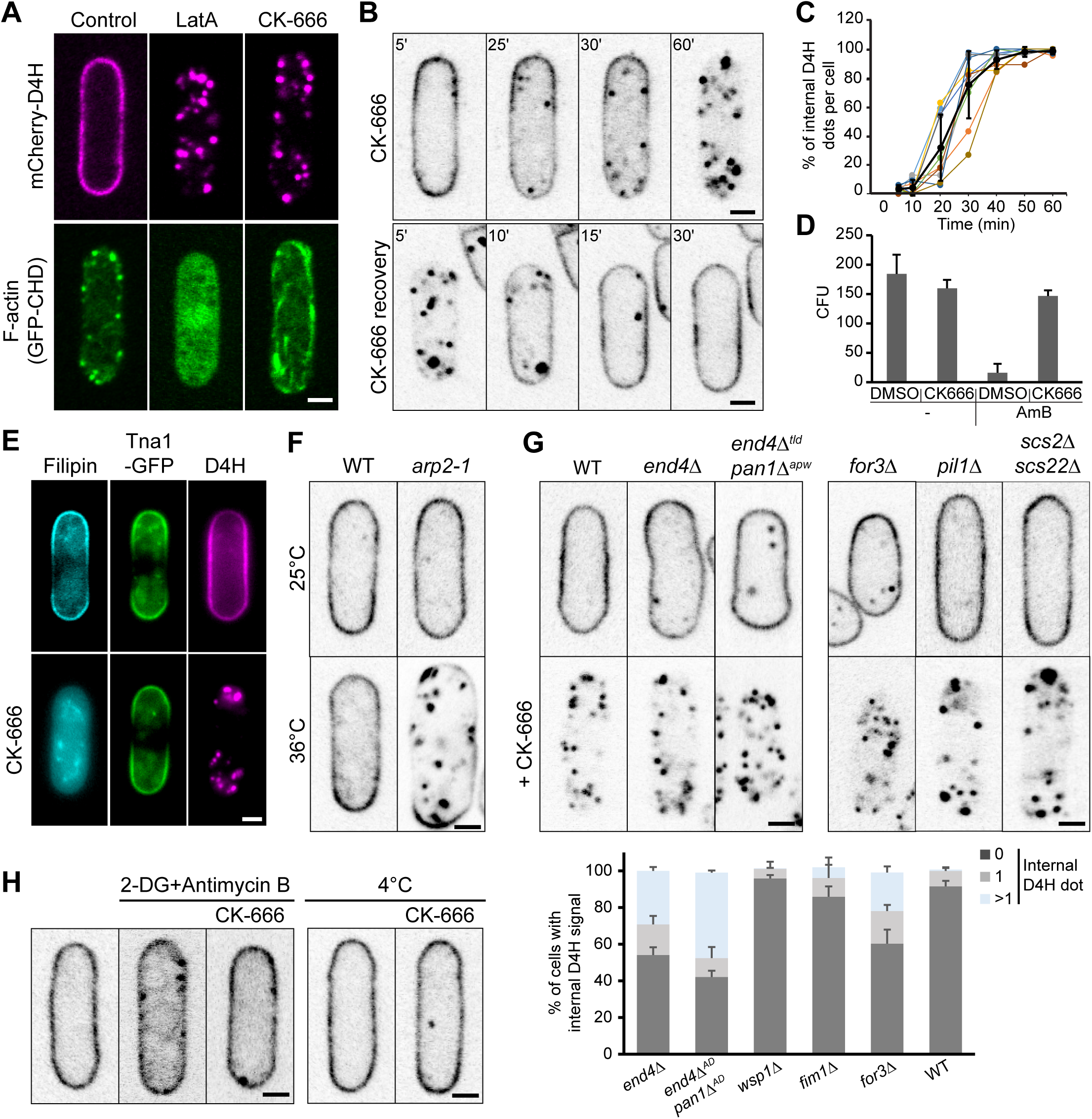
Internalization of sterols upon inhibition of Arp2/3 activity. **A.** Localization of mCherry-D4H and F-actin marker GFP-CHD upon depolymerization of F-actin by latrunculin A (LatA, 200μM) or Arp2/3 inhibition by CK-666 (500μM). **B.** Time-lapse sequence of mCherry-D4H after CK-666 addition (upper panel) and wash-out after 1h treatment (lower panel) **C.** Quantification of the number of internal D4H-positive structures in CK-666-treated cells imaged every 5 min. Values are normalized to the maximal number of internal D4H dots in the z-stack. **D.** Wildtype cells become resistant to Amphotericin B (AmB) upon CK-666 treatment. The graph shows the viability of cells pretreated with CK-666 (or DMSO as control) for 1h and then treated with a high dose (4μg/ml) of AmB for another hour. N = 3 experiments. Error bars show the standard deviation. **E.** Loss of filipin staining, but not PM GFP-Tna1 after 1h CK-666 treatment. **F.** mCherry-D4H in *arp2-1* mutant cells grown at 25°C or 36°C for 6h. **G.** mCherry-D4H in selected endocytic (*end4Δ*, *end4*^*Δtld*^ *pan1*^*Δapw*^), actin cable (*for3Δ*), eisosome (*pil1Δ*) and ER-PM attachment (*scs2Δ scs22Δ*) mutants treated with CK-666 for 1h. The bottom graph displays the number internal dots in z-stacks (n ≥ 30 cells per strain). **H.** mCherry-D4H relocalization upon energy depletion. (Left) Cells were treated for 1h with deoxyglucose (DG) and antimycin to deplete energy, and then incubated for 1h more with CK-666. (Right) Cells were precooled on ice for 30 min and then treated with CK-666 for 1h on ice. Scale bars 2μm.

Because Arp2/3 inhibition blocks clathrin-mediated endocytosis in yeast, we probed whether other means of endocytosis inhibition would also provoke sterol internalization (Fig 2G). As endocytosis is essential, we used a range of viable mutants of variable severity and quantified the number of internal D4H-positive structures. Deletion of *wsp1* or *fim1*, which moderately affect endocytic patch dynamics, had no effect (Lee et al., 2000; Sirotkin et al., 2005; Skau and Kovar, 2010). Even cells with severely impaired endocytosis, lacking the adaptor protein End4 or carrying End4/Pan1 alleles unable to bind actin (*end4*^*Δtld*^ *pan1*^*Δapw*^) (Chen and Pollard, 2013; Iwaki et al., 2004), retained D4H at the PM and showed only moderate numbers of internal D4H-positive structures. Disrupting Arp2/3 in these mutants led to D4H internalization as in wildtype cells. Thus, sterol internalization is not caused by a block in clathrin-mediated endocytosis, but by inhibition of another Arp2/3 function.

We entertained the possibility that sterols may be internalized through a non-canonical actin-independent endocytic pathway. However, deleting the homologues of genes involved in non-clathrin-mediated endocytosis in mammalian cells (Doherty and McMahon, 2009), such as dynamin (*vps1*) or *arf6* did not block CK-666-induced D4H internalization (Fig S3). We were also unable to identify a resident transmembrane protein that would co-internalize with D4H: from a screen of over 24 PM transmembrane proteins, none co-internalized with D4H, though some decorated internal structures in addition to the PM even in untreated conditions (Fig S4A-B). Notably, Tna1 did not co-internalize with D4H (Fig 2E). The lipophilic dye FM4-64 also did not internalize in CK-666-pretreated cells (Fig S4C). Internalization was specific to sterol lipids, as neither a GFP-2xPH(PLC∂) nor a GFP-LactC2 probe, which bind PIP_2_ and PS respectively, internalized like D4H upon CK-666 treatment (Fig S4D). We conclude that sterol internalization from the PM is unlikely to involve direct membrane endocytosis. We also probed for a possible role of eisosomes, plasma membrane invaginations present in fungi and other walled eukaryotes, with suggested enrichment in sterol (Grossmann et al., 2007; Kabeche et al., 2015b). However, deletion of the major eisosome structural component Pil1 did not block CK-666-induced D4H internalization (Fig 2G) (Kabeche et al., 2011). The process was also independent of ER-PM contacts mediated by the VAP-family tethers Scs2 and Scs22 (Fig 2G) (Zhang et al., 2012). Interestingly, D4H translocation is an active process that requires energy since blocking ATP production with antimycin A or incubation of cells at 4°C blocked CK-666-induced D4H internalization (Fig 2H). In summary, sterol internalization upon Arp2/3 inhibition is an endocytosis- and eisosome-independent active process.

### Sterols accumulate within the endosomal compartment upon Arp2/3 inhibition

We examined the nature of the internal D4H-positive structures after Arp2/3 inhibition by correlative light-electron microscopy (CLEM). mCherry-D4H expressing cells were treated for 45 min with CK-666 and immediately cryo-fixed by high pressure freezing. After freeze substitution, 300-nm sections were imaged by both fluorescence and electron microscopy. We focused on cells that contained internal and PM signal, with the latter used to co-align the light and electron microscopy images. The vast majority of D4H internal signals (144 of 154 foci in 44 cells) corresponded to membrane-enclosed compartments, which we name <u>st</u>erol-<u>ri</u>ch <u>c</u>ompartments (STRIC). 77.3% of these corresponded to large, often electron-dense, almost spherical organelles with a diameter of 128.5 ± 26.9 nm (Fig 3A). The other signals overlapped with larger membrane organelles (18.7%) (Fig 3B), possibly Golgi (see below), or PM invaginations (4%). We acquired tilt-series and reconstructed tomograms for 104 STRIC, which confirmed their vesicular nature (Fig 3A-B).

**Figure 3.**
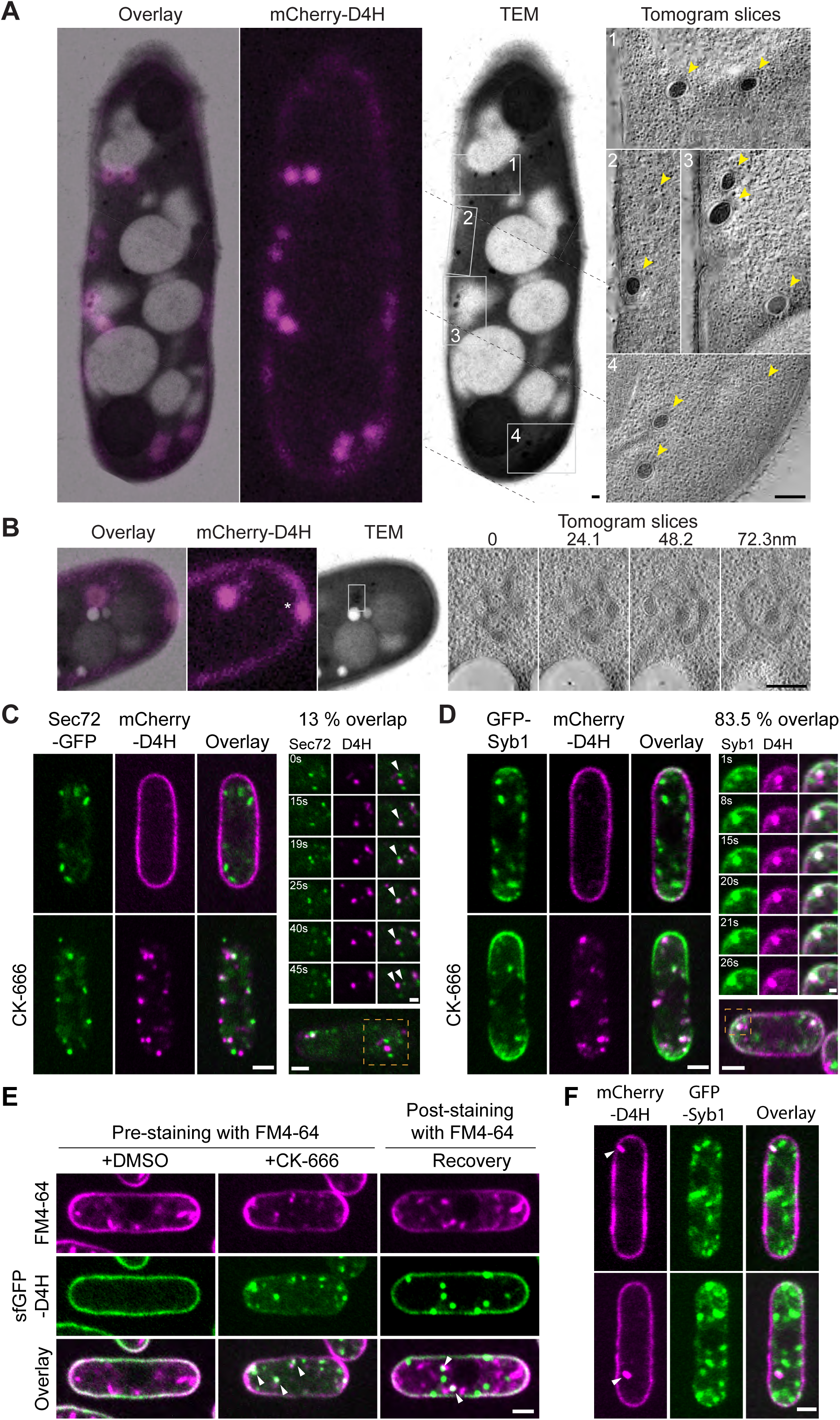
Sterols are internalized to endosomes. **A-B.** Correlative transmission electron microscopy (TEM) and epifluorescence (mCherry-D4H) images of a 300nm section of mCherry-D4H-expressing cells treated with CK-666 for 45 min-1h. The overlay is shown on the left. In (A), the TEM image is a composite of three micrographs (junctions indicated by dashed lines). Virtual sections through tomographic reconstruction of the four regions highlighted by grey boxes are shown on the right. In (B), serial virtual sections at 24.1 nm distance through tomographic reconstruction of the boxed region is shown on the right. Yellow arrowheads point to D4H-positive spherical organelles. **C-D.** Colocalization of mCherry-D4H and Sec72-GFP (C) or GFP-Syb1 (D) in cells with or without CK-666 for 1h. An example time-lapse sequence is shown on the right. 17% of D4H internal dots overlapped with Sec72-GFP and 84.7% with GFP-Syb1 upon CK-666 treatment (n = 20 cells). **E.** Colocalization of internal D4H structures with internalized FM4-64 in cells prelabelled with FM4-64 and treated 30min with CK-666 (left) or labelled with FM4-64 upon CK-666 wash-out (right). **F.** Untreated wildtype interphase cells with internal mCherry-D4H signal, which colocalizes with Syb1. Scale bars are 200nm in (A-B), 1μm in (D, right panel), 0.5μm in (C, right panel) and 2μm elsewhere.

To examine the STRIC identity, we labelled CK-666-treated cells expressing mCherry-D4H with a large panel of organellar markers. We did not detect significant colocalization with markers labelling ER exit sites, early Golgi, pre-vacuolar compartment, endocytic patches, or post-Golgi vesicles (Fig S5). STRIC also did not colocalize with lipid droplets or mitochondria (Fig S5). By contrast, we detected occasional colocalization with the late Golgi/early endosome marker Sec72-GFP (Fig 3C): in time-lapse experiments, initially distinct D4H and Sec72 signals transiently overlapped for several time frames before dissociating again (Fig 3C, Movie S4). This suggests that STRIC transiently associates with late secretory compartments, consistent with the CLEM data. Importantly, D4H strongly colocalized with the v-SNARE Synaptobrevin-like Syb1, a core component of the vesicle-PM fusion machinery, which resides on vesicles and is recycled from the PM (Fig 3D, Movie S5). In untreated WT cells, Syb1 localizes to punctate structures that correspond to endosomal compartments (Edamatsu and Toyoshima, 2003; Gachet and Hyams, 2005), as shown by strong colocalization with the lipophilic dye FM4-64 internalized for 5 min (Fig S6). A fraction of Syb1 also localizes to the PM, where it is trapped upon endocytosis arrest by CK-666 treatment. In these conditions, D4H colocalized with the residual internal Syb1-GFP signal. STRIC could also be labelled by uptake of FM4-64 if applied before CK-666 treatment or at the time of CK-666 wash-out (Fig 3E). Together our findings indicate that blocking Arp2/3 activity results in accumulation of sterols within the endosomal compartment. As D4H binds only to membranes containing more that 20mol% of sterol, this indicates enormous accumulation of sterols in endosomes.

Because internal D4H dots were also observed in untreated wildtype cells (Fig 1A, Movie S1; 20.8 ± 2.5% of all cells), we asked whether these also correspond to STRIC labelled by Syb1-GFP. Indeed, 32.6 ± 6% of these naturally-forming D4H internal dots coincided with Syb1-positive structures (Fig 3F). These observations suggest that STRIC also form in unperturbed wildtype cells, but are enriched upon Arp2/3 inhibition.

### Sterols travel between the PM and endosomes independently of the secretory pathway

To assess the importance of the secretory pathway in sterol internalization to endosomes, we treated cells with Brefeldin A (BFA). BFA is a macrocyclic lactone that inhibits the Sec7-family of ARF exchange factors, leading to collapse of the Golgi compartment (Peyroche et al., 1999). In *S. pombe*, BFA blocks secretion and leads to Golgi disappearance with concomitant ER accumulation (Turi et al., 1994). Indeed, BFA caused the redistribution of the early Golgi marker Anp1-GFP to the ER (Fig 4A). Moreover, it induced the dispersion of most Syb1 signal, indicating loss of most or all ER-anterograde trafficking (Fig 4B). 1h BFA treatment had no effect on D4H localization (Fig 4A). However, consistent with sterol trafficking through endosomes, in cells pre-treated with BFA for 30 min to induce Syb1 dispersion, Arp2/3 inhibition largely failed to induce the formation of STRIC: the D4H signal either remained at the PM (Fig 4Bi) and/or showed an internal haze and/or vacuolar staining (Fig 4Bii, iii). Interestingly, occasional internal dots (Fig 4B, i and ii) coincided with remaining Syb1-positive structures. These data indicate that sterol internalization from the PM to endosomes is strongly perturbed upon BFA treatment, with re-direction of sterols to other endo-membranes and residual transport to remaining Syb1-positive structures.

**Figure 4.**
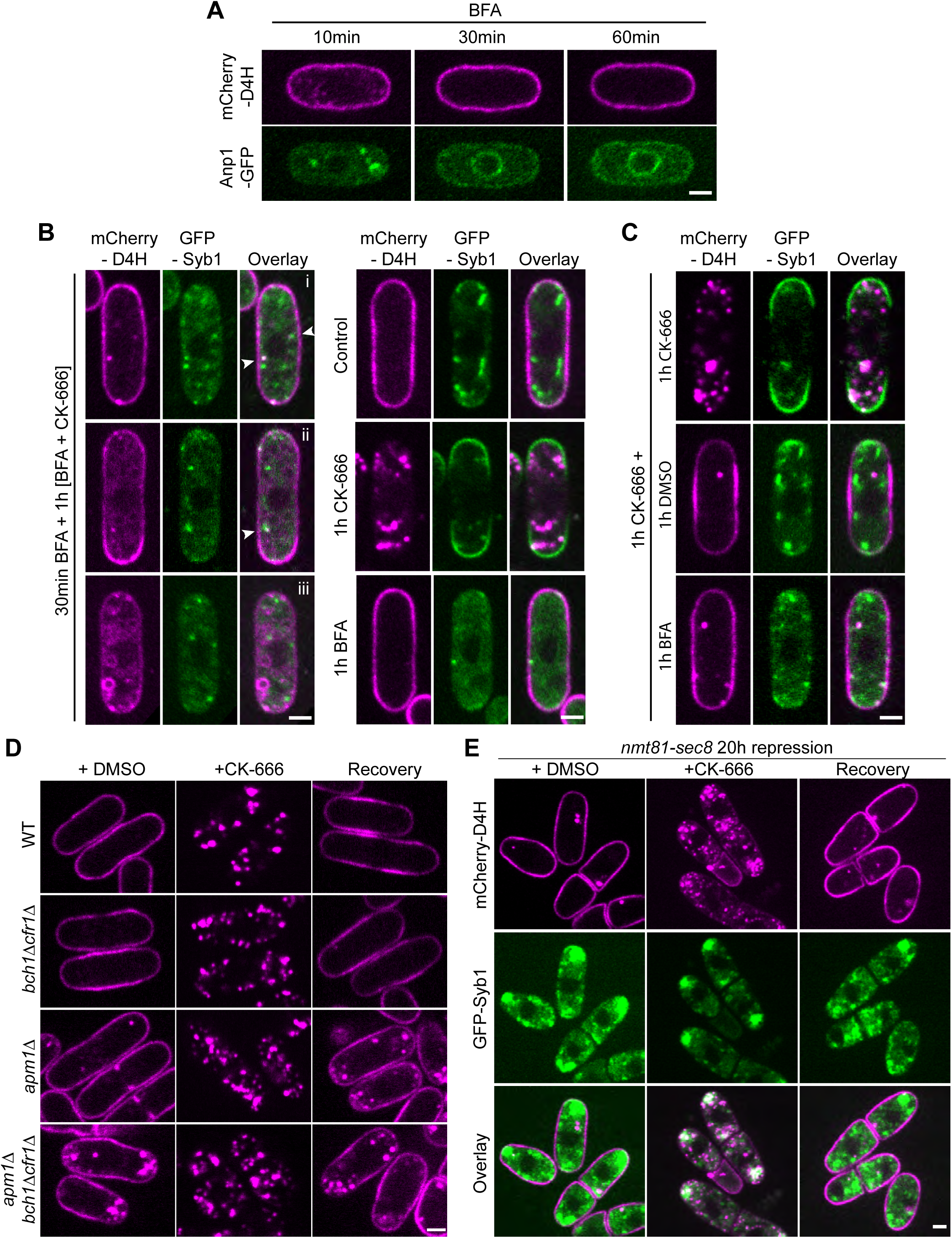
Sterol transport is independent of vesicle trafficking. **A.** Treatment of cells with 300μM BFA leads to Golgi, marked by Anp1-GFP, collapse to the ER, but has no effect on D4H distribution at the PM. **B.** Cells pretreated with BFA for 30 min to induce Golgi collapse and then incubated in the presence of BFA and CK-666 to induce sterol internalization display three principal phenotypes: i) D4H at the PM with occasional internal dot-like structure that coincided with Syb1 (arrowheads), ii) D4H at the PM and internal haze, with dot like-structures that coincided with Syb1, iii) weak D4H at the PM with internal haze and vacuolar staining. The right panel shows control treatments. **C.** Cells were treated for 1h with CK-666, and then 1h more with CK-666, DMSO or BFA. **D.** mCherry-D4H localization during steady-state growth (DMSO), upon 1h CK-666 treatment, and after 1h recovery in exomer deficient mutants (*bch1Δ cfr1Δ*), mutants lacking the AP-1 adaptor complex subunit (*apm1Δ)* and triple mutant (*bch1Δ cfr1Δ apm1Δ)*. **E)** mCherry-D4H localization during steady-state growth (DMSO), upon 1h CK-666 treatment, and after 1h recovery in cells in which *sec8* expression (under the control of the *p*^*nmt81*^ promoter) has been repressed for 20h. Scale bars 2μm

Restoration of Arp2/3 function upon CK-666 wash-out leads to D4H return to the PM (Fig 2B, Movie S3). Interestingly, addition of BFA at the time of CK-666 removal did not impair D4H signal recovery to the PM (Fig 4C). Previous work showed that trafficking from the endosome is regulated by the clathrin adaptor protein Apm1 (AP-1), which localizes to endosomes, and *apm1Δ* accumulates post-Golgi vesicles (Kita et al., 2004). The exomer complex, formed by Bch1 and Cfr1, which also localizes to endosomes, plays a more minor role, though *apm1Δ exomerΔ* mutants display enhanced defect in trafficking from the endosome (Hoya et al., 2017). Consistent with aberrant endosome organization, *apm1Δ* and *apm1Δ exomerΔ* triple mutants showed extensive D4H internal signal (Fig 4D). However, treatment of these cells with CK-666 led to D4H internalization as in wildtype cells, with recovery of D4H to the PM upon CK-666 wash-out to levels observed during steady-state growth (Fig 4D). The exocyst complex, which promotes the tethering of secretory vesicles to the PM, is critical for polarized secretion (Heider and Munson, 2012), and depletion of the essential exocyst subunit Sec8 leads to the sub-apical accumulation of Syb1-positive organelles (Bendezu and Martin, 2011). Sec8 depletion had no substantial effect on D4H distribution, which remained at the PM (Fig 4E). Again, treatment of these cells with CK-666 led to D4H internalization to Syb1 compartments as in wildtype cells and CK-666 wash-out permitted full recovery to the PM, despite the sub-apical retention of Syb1 (Fig 4E). We conclude that, similar to retrograde transport from PM to endosomes, anterograde transport of sterols from the endosome to the PM is independent of the canonical vesicular trafficking machinery.

### Sterol distribution in sterol transport mutants

Sterol trafficking may involve sterol-binding proteins that transport sterols between membranes. Studies in mammalian and budding yeast cells have implicated the ORP (OSBP Related Proteins) and STARkin/LAM (StART-like) protein families in this process (Luo et al., 2019). *S. pombe* encodes 6 ORP and 2 STARkin proteins, which we named according to phylogenetic relationship with the *S. cerevisiae* homologues (Fig 5A). To probe the possible role of these proteins in sterol transport, we generated gene deletions and analysed the distribution of mCherry-D4H in these strains (Fig 5B). In most of the mutants D4H localized correctly to the PM under standard growth conditions, with two exceptions. First, cells lacking Osh41 showed only a weak PM signal, and a significant fraction of D4H localized in the cytosol and on small internal structures (Fig 5B, left panel). Second, cells lacking Ltc1 showed D4H at the PM, but the signal was significantly depleted from cell poles (Fig 5B, left panel; Fig 5C, Movie S6).

**Figure 5.**
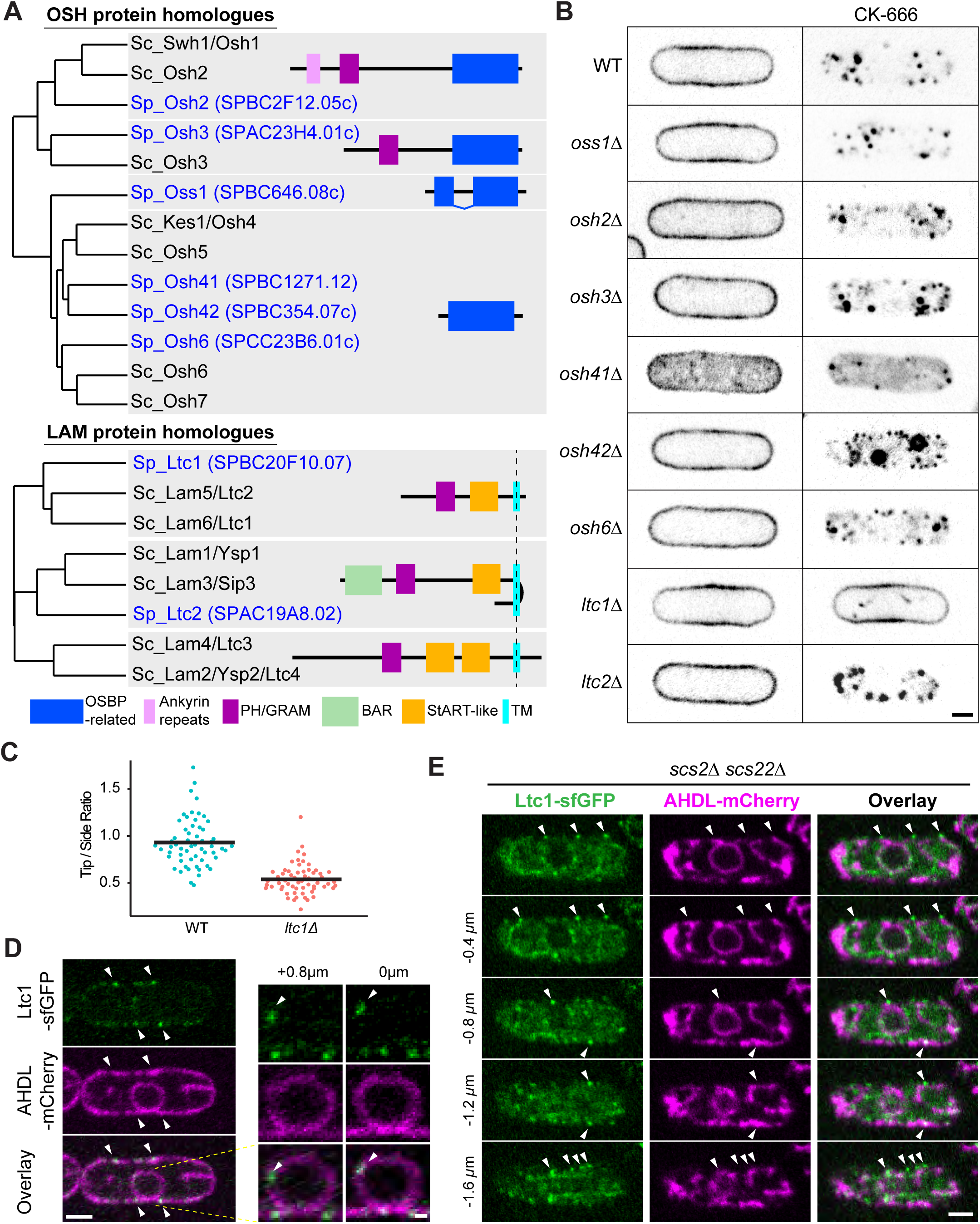
Ltc1 is required for sterol internalization and localization to ER-PM contact sites. **A.** Neighbour-joining phylogenetic trees of *S. pombe* and *S. cerevisiae* OSH and LAM-family proteins. The *S. pombe* genes were named in analogy to the *S. cerevisiae* names, with Osh = <u>O</u>xy<u>S</u>terol-binding protein <u>H</u>omologue, Oss1 = <u>O</u>xy<u>S</u>terol binding <u>S</u>plit-domain protein and Ltc1/2 = <u>L</u>ipid <u>T</u>ransfer at <u>C</u>ontact site. Protein domains identified by SMART are indicated. **B.** mCherry-D4H in OSH-family or LAM-family deletion mutants treated or not with CK-666 for 1h. **C.** Quantification of mCherry-D4H fluorescence intensity at cell tip versus cell side. Bars show the mean (n ≥ 50 cells). **D.** Ltc1-GFP expressed from the native genomic locus localizes to cortical and internal dots (arrowheads) at the ER, labelled with mCherry-AHDL. Right panels show an enlargement of the nucleus. **E.** Ltc1-GFP localizes to residual ER-PM contacts (arrowheads) in the *scs2Δ scs22Δ* mutant. Panel represents middle plane view and 4 subsequent planes at 0.4μm distance. Scale bars are 0.5μm in (D, right panel) and 2μm elsewhere.

We tested the ability of the deletion mutants to internalize PM sterols to endosomes upon CK-666 treatment. Again, most mutants were proficient in sterol internalization to endosomes, including cells lacking Osh41, with two exceptions. First, *osh42Δ* showed re-localization not only to endosomes but also to vacuoles (Fig 5B, right panel). Second, *ltc1Δ* completely blocked endosomal sterol enrichment and D4H remained PM-associated in this mutant (Figure 5B, right panel; Movie S7). We thus further focused on deciphering the role of Ltc1 in the process of sterol accumulation in endosomes.

### Ltc1 localizes to ER-PM contact sites and regulates PM to endosome sterol flow

To probe the localization of Ltc1, we constructed a C-terminally sfGFP-tagged allele expressed from the native genomic locus. Ltc1-sfGFP formed weak punctate structures principally at the cell cortex and occasionally in the cell interior including the perinuclear ER (Fig 5D). As the native signal was very weak we expressed Ltc1-sfGFP under the control of the strong actin (*p*^*act1*^) or weaker pom1 (*p*^*pom1*^) promoter in *ltc1Δ* cells. Both constructs rescued the ability of Ltc1 protein to promote endosomal sterol enrichment upon CK-666 treatment (Fig 6A). Strong protein overexpression from *p*^*act1*^ led to labelling of the entire ER, indicating that Ltc1 is an ER-resident protein. Expression from *p*^*pom1*^ produced similar, but stronger, punctate pattern as from the native locus (Fig 6A). Interestingly, in *scs2Δ scs22Δ* double mutants that exhibit fewer ER-PM contacts than wildtype cells, cortical Ltc1-sfGFP puncta (expressed from the native locus) were observed at these residual contact sites (Fig 5E), indicating Ltc1 marks VAP-family-independent ER-PM contact sites. We conclude that the majority of Ltc1 is enriched at ER-PM contact sites.

**Figure 6.**
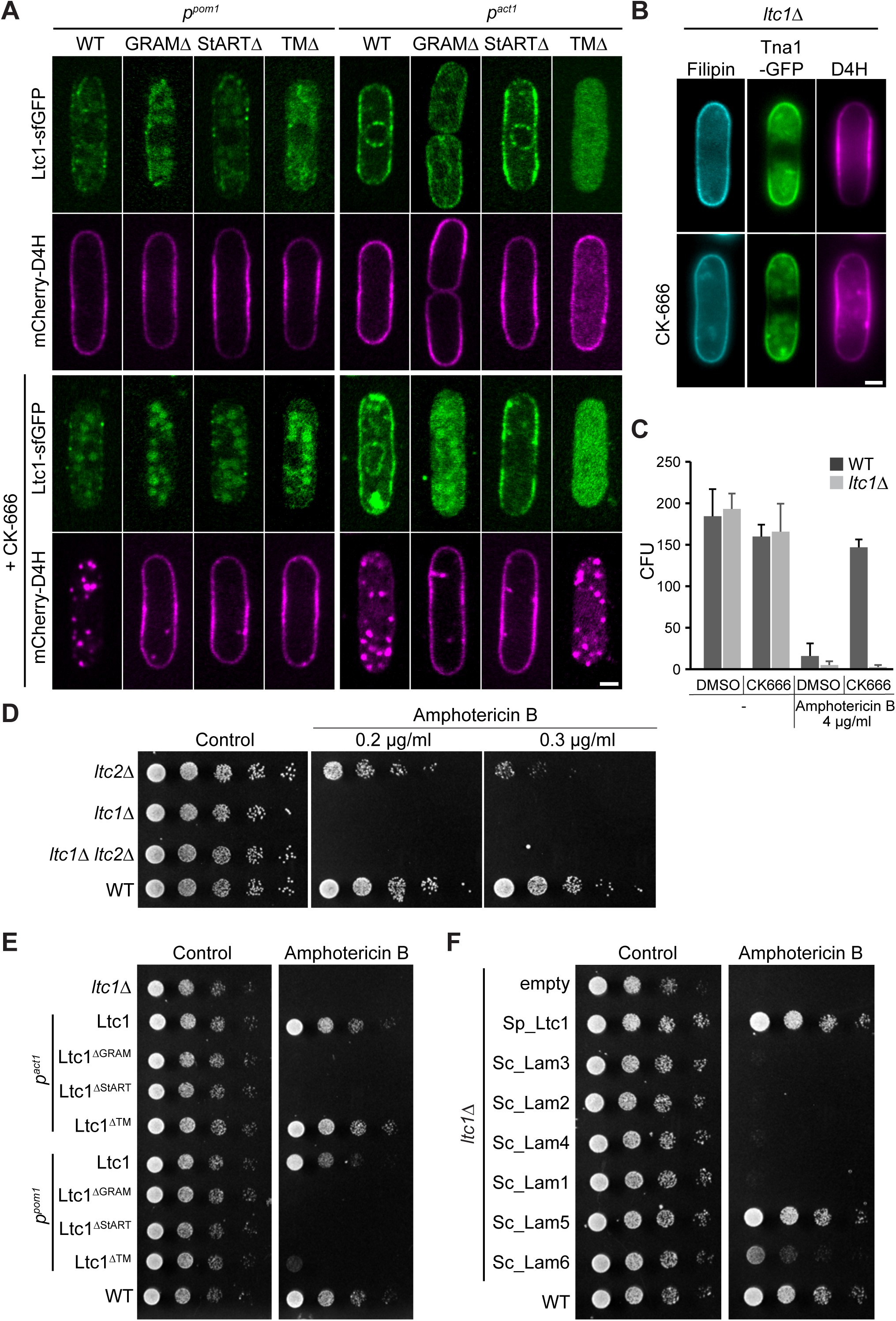
Ltc1 StART-like domain is essential for retrograde sterol transfer. **A.** Ltc1-sfGFP and truncated alleles lacking GRAM, StART-like or TM domain expressed under control of the weak *p*^*pom1*^ promoter or strong *p*^*act1*^ promoter in *ltc1Δ* cells, treated or not with CK-666 for 1h. **B.** Filipin remains at the PM of *ltc1Δ* cells treated with CK-666 for 1h. **C.** WT but not *ltc1Δ* cells become resistant to Amphotericin B (AmB) during CK-666 treatment. WT data and methodology as in Fig 2D. **D**. Serial dilutions of WT and indicated mutants on EMM-ALU media containing AmB. **E.** Serial dilutions of *ltc1Δ* strains expressing WT Ltc1 or truncated alleles under the control of *p*^*pom1*^ and *p*^*act1*^ on plates containing 0.2μg/ml AmB. **F.** Serial dilutions of *ltc1Δ* mutant expressing *S. cerevisiae* LAM proteins under the control of *p*^*act1*^ on plate containing 0.2μg/ml AmB. Scale bars 2μm.

The Ltc1 protein sequence exhibits three notable domains: a PH-like GRAM domain, a StART-like domain, predicted to bind sterols, and a C-terminal transmembrane domain (Fig 5A). To determine the importance of these domains for Ltc1 localization and function, we expressed mutants lacking these domains, (Ltc1^ΔGRAM^, Ltc1^ΔStART-like^ and Ltc1^ΔTM^) under the control of *p*^*act1*^ or *p*^*pom1*^ promoter (Fig 6A). Ltc1^ΔGRAM^ was expressed very poorly even from *p*^*act1*^ and only in few cells could we detect an ER signal. Under *p*^*pom1*^ control, expression levels were too low to test its presence at ER-PM contacts. Ltc1^ΔStART-like^ displayed a proper punctate localization pattern and was expressed at levels similar to WT. Ltc1^ΔTM^ localized to the cytosol. At low expression levels, none of these mutants was able to complement the *ltc1Δ* phenotype (Fig 6A, left panel). However, upon overexpression, the *ltc1*^*ΔTM*^ allele promoted an increase in cytosolic D4H in untreated cells and rescued the ability to internalize sterols to endosomes in presence of CK-666 (Fig 6A, right panel). In summary, these data are consistent with the view that Ltc1 promotes sterol transport through its essential StART-like domain and that the main function of the TM domain is to increase the concentration of Ltc1 at sites of sterol transport.

### Sterol flow contributes to plasma membrane homeostasis

In line with the retention of D4H at the PM in *ltc1Δ*, these cells retained PM filipin staining and, in contrast to WT cells, did not become resistant to high concentrations of AmB after treatment with CK-666 (Fig 6B-C). Cells lacking *ltc1* showed enhanced AmB sensitivity even in steady-state conditions (Fig 6D). As for D4H internalization, AmB sensitivity was complemented by overexpression of Ltc1^ΔTM^, but not any other truncation mutant (Fig 6E). We used this drug sensitivity to test for functional homologues amongst LAM-family proteins, which showed that both *S. cerevisiae* Lam5 and Lam6 were able to restore AmB resistance to *ltc1Δ* cells (Fig 6F). This finding is consistent with phylogenetic proximity of Ltc1 and these two *S. cerevisiae* LAM-family proteins (Fig 5A). Together these results indicate that *ltc1Δ* has excess sterols at the PM not only by failing to internalize sterols upon CK-666 treatment, but also during steady state growth.

Remarkably, *ltc1Δ* cells not only did not internalize sterols but formed long PM invaginations upon CK-666 treatment. Residual internal D4H-mCherry structures observed in CK-666-treated *ltc1Δ* cells were not vesicular as in WT cells but were elongated and connected to the PM, as shown by confocal sectioning. These PM invaginations were decorated by not only D4H-mCherry, but also resident PM proteins, such as the t-SNARE Psy1 (Fig 7A). CLEM of D4H-mCherry in CK-666-treated *ltc1Δ* cells showed that out of 32 internal fluorescence signals, 22 corresponded to membrane tubes or sheets (Fig 7B, i and iii), occasionally forming larger bulges or more complex structures (Fig 7Bii). In 4 cases, connection of the tube to the PM was captured within the section (Fig 7Bi). In several instances, these tubes were in very close proximity to ER structures (Fig 7Biii). We also observed 2 signals overlapping with grooves at the PM. We did not observe any overlap between the D4H signal and large electron-dense vesicles as in WT cells, though 2 signals overlapped with smaller vesicles. Interestingly, we also found close contact between D4H-labelled invaginations and AHDL-labelled ER by light microscopy (Fig 7C). We conclude that blocking both endocytosis and sterol flow to endosomes leads to an excess of PM, which now forms extended ER-proximal internal invaginations.

**Figure 7.**
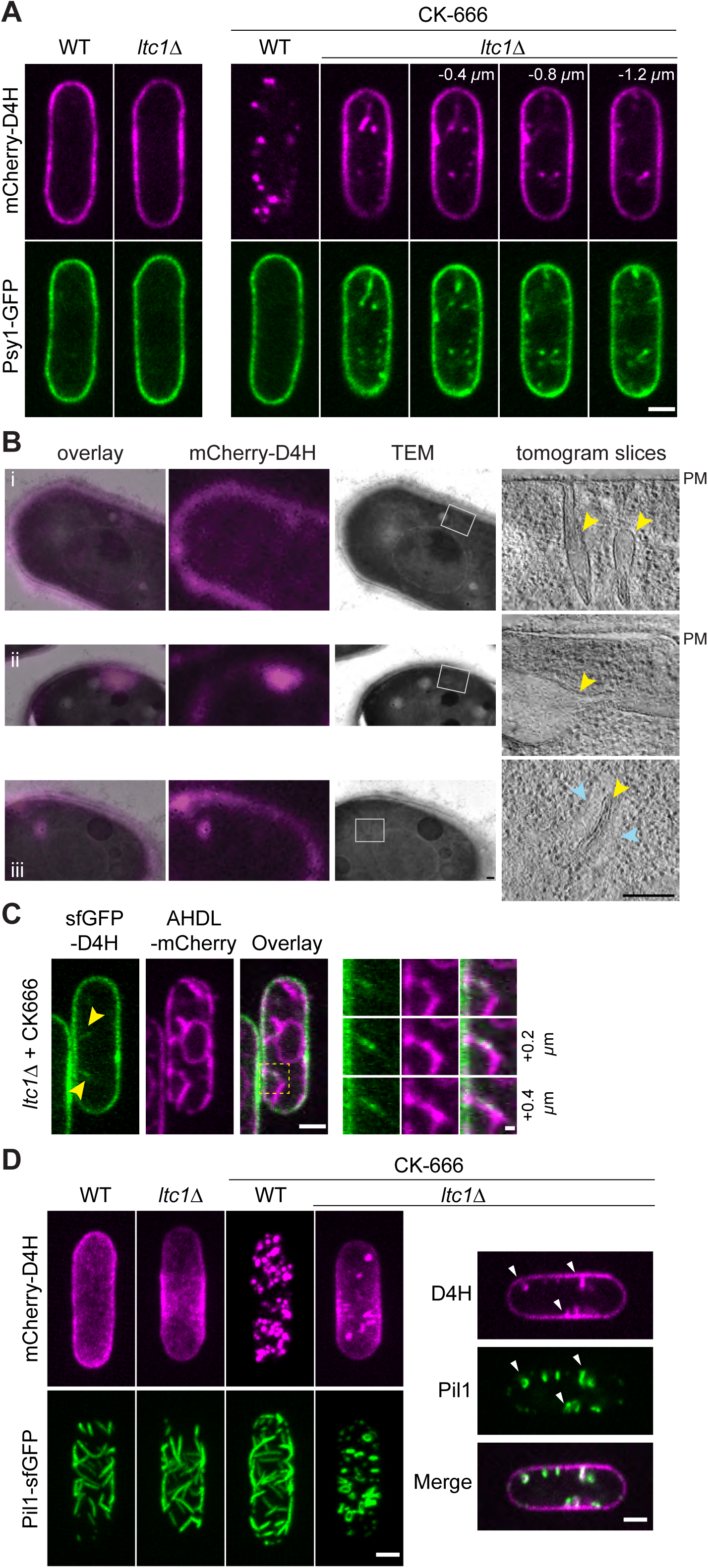
Deficiency in retrograde sterol transfer leads to large PM invaginations. **A.** Internal mCherry-D4H signal in CK-666-treated *ltc1Δ* cells form deep invaginations co-labelled by the PM tSNARE GFP-Psy1. **B.** Correlative transmission electron microscopy (TEM) and epifluorescence (mCherry-D4H) images of a 300nm section of mCherry-D4H-expressing *ltc1Δ* cells treated with CK-666 for 45 min-1h. The overlay is shown on the left. Virtual sections of tomographic reconstruction of the regions highlighted by grey boxes are shown on the right. Three examples show (i) invaginations from the PM, (ii) large internal bulging region, and (iii) tube surrounded by ER (blue arrowheads). Yellow arrowheads point to D4H-positive organelles. **C.** PM invaginations labelled by sfGFP-D4H in CK-666 treated *ltc1Δ* mutant are surrounded by ER marked by mCherry-AHDL. An enlargement of the boxed region is shown over three z-stacks on the right. **D.** PM invaginations labelled by sfGFP-D4H in CK-666 treated *ltc1Δ* form at eisosomes, marked by Pil1. In WT cells, Pil1 and D4H do not colocalize. In *ltc1Δ* cells, Pil1 appears to surround the D4H signal on PM invaginations. Images on the left are maximum projections of 0.4μm-spaced z-planes. Scale bars are 200nm in (B), 0.5μm in (c, righ panel) and 2μm elsewhere.

The invaginations were marked by Pil1, suggesting that they arise from eisosome expansion (Fig 7D). Pil1-marked eisosomes appeared normal in *ltc1Δ*, forming extended furrows in the plane of the membrane, consistent with their described organization in WT cells. However, addition of CK-666 to *ltc1Δ* cells caused a massive reorganization of the Pil1-marked structures, which lost their planar organization and instead protruded towards the cell interior (Fig 7D). By contrast, addition of CK-666 to wildtype cells caused a much more modest reorganization of eisosomes, which now also formed at the poles of the cell, likely due to growth arrest (Fig 7D). Together, these data show that simultaneous block of endocytosis and sterol internalization from the PM leads to excess PM, which leads to aberrant enlargement and alteration of eisosomes.

## Discussion

In this study, we have used a genetically-encoded fluorescent probe, D4H, to detect sterol-rich membranes in live cells, which revealed an intracellular sterol flux: sterols are carried from the PM to endosomes, likely through the ER, with the help of the lipid transfer protein Ltc1, and back to the PM independently of classical vesicular transport.

### The D4H biosensor gives novel insights into sterol distribution in cellular membranes

Because D4H is genetically encoded and non-toxic, it makes a highly valuable sterol biosensor. In unperturbed cells it primarily labels the PM, but also occasionally intracellular TGN/endosomes. Surprisingly, the sterol distribution detected by D4H at the PM is distinct from that of filipin, which has been widely used to describe sterol-rich domains in fungi. D4H is fairly homogeneous along the PM or even depleted at growth sites, which contrasts with the labelling of cell poles and division sites by filipin. Because filipin is applied from the cell outside and D4H is expressed inside, this discrepancy may suggest distinct sterols distributions in the outer and inner PM leaflets, with higher concentration of sterols in the outer leaflet at sites of polarized growth. Indeed, other approaches have also suggested an asymmetry in sterol distribution between the two yeast PM leaflets (Solanko et al., 2018). Alternatively, filipin and D4H localization may be influenced by differences in sterol accessibility, for instance due to complex formation with other lipids (e.g. sphingolipids). The distinct filipin staining pattern may also reflect cell-wall permeability since both tips and division sites are places of strong cell-wall remodeling. A clear advantage of D4H is its ability to non-invasively track sterol-rich endo-membranes (containing >20 mol%) in live cells, which filipin’s toxicity does not permit. We note that the transmembrane protein Tna1, which was previously proposed to label sterol-rich domains (Makushok et al., 2016), also fails to track sterol on endo-membranes.

D4H revealed alterations in sterol distribution for the deletion of three predicted sterol-binding LTPs: single deletions of *osh41*, *osh42* and *ltc1* all affected D4H localization, though in distinct ways. While we have focused here on the role of Ltc1 in retrograde sterol flow from the PM, our observations suggest that Osh41 may promote PM sterol levels during steady-state growth and Osh42 may control sterol flow away from vacuoles. Osh41 and Osh42 are related to *S. cerevisiae* Osh4, which binds sterols *in vitro* (de Saint-Jean et al., 2011; Im et al., 2005; Schulz et al., 2009) and is proposed to promote ER to PM sterol transport by using the counter-gradient of PI4P (Moser von Filseck et al., 2015), although *in vivo* data suggest a role in membrane sterol organization rather than sterol transport (Georgiev et al., 2011; Quon et al., 2018; Raychaudhuri et al., 2006). These observations demonstrate that D4H is a highly valuable tool to study the intracellular trafficking of sterols.

### Retrograde trafficking of sterols from the plasma membrane to endosomes

We discovered that inhibition of Arp2/3-mediated actin assembly triggers the internalization of D4H from the PM to intracellular sterol-rich compartments (STRIC). These organelles are labeled by late Golgi/endosome markers, such as the Arf GEF Sec72 and the v-SNARE Synaptobrevin-like Syb1, as well as by the internalized FM4-64 dye. The morphology and electron density of the STRIC is in fact highly reminiscent of that of secretory vesicles, suggesting these may represent a recycling/sorting compartment destined to the PM. The overlay of D4H and Syb1 signals in exocyst-depleted cells, which accumulate 100nm-diameter vesicles (Wang et al., 2002), further supports this view. Thus, D4H is relocalized to endosomes. Endosomal sterol enrichment occurs beyond fungi. In mammalian cells, the endosomal/lysosomal compartment plays an important role during sterol re-distribution to the PM after uptake of LDL-derived cholesterol (Kanerva et al., 2013). In plant cells, sterols were also shown to accumulate in endosomes upon disruption of the actin cytoskeleton (Grebe et al., 2003). Thus, actin-dependent endosome-PM sterol exchange may be a conserved feature of sterol trafficking.

Importantly, the sterol-rich endosomes are distinct from early endocytic vesicles. Four lines of evidence show that sterols do not travel from the PM to endosomes in endocytic vesicles. First, massive sterol internalization occurs after inhibition of Arp2/3 with CK-666, which blocks clathrin-mediated endocytosis in yeast (Gachet and Hyams, 2005; Galletta and Cooper, 2009) Second, internalization occurs normally in cells in which endocytosis is genetically impaired. Third, despite an extensive screen we did not identify any PM marker, whether peripherally associated or transmembrane, that co-internalizes with D4H. Fourth, sterol internalization is not affected by mutants predicted to affect membrane sculpting (BAR protein or dynamin mutants), but is blocked in absence of the previously uncharacterized LAM-family protein Ltc1. We conclude that sterols do not reach the endosome through an endocytic route, but are transported between membranes by lipid transfer proteins.

What route do sterols follow from the PM to endosomes? Because D4H labels the PM and endosomes, but no other endo-membrane, and because sterol depletion from the PM is inefficient in absence of the Syb1-compartment, the simplest explanation would be a direct transport from PM to endosomes. However, several considerations indicate that the transport route is less direct. First, Ltc1 is an ER resident that localizes to ER-PM contact sites, suggesting that it promotes sterol transport between these two organelles. Second, in *ltc1Δ*, we observed tight ER contacts with expanded regions of the PM, suggesting that these contacts exist independently of successful sterol transport. Thus, a likely route involves sterols flowing from the PM to the ER and then from the ER to endosomes. An estimate of the number of sterol molecules internalized and the space on endosomes also reveals that not all sterol molecules internalized from the PM may have space on the endosomes: Let’s consider a 10μm-long yeast cell internalizing sterols to 50 endosomes (upper bound of our observations). While the surface of the PM is about 125μm^2^, the total endosomal surface is only about 2.5μm^2^, i.e. 50-fold lower. If PM sterol levels drop from a steady-state concentration of 30 mol% (detectable by both D4 and D4H) to just below 20 mol% (just below D4H detection levels), and endosomes accommodate a massive 50 mol% sterol concentration increase, this would only account for 10% of the moved sterols. This suggests that a large pool of internalized sterols is also present at other locations than endosomes. We hypothesize that these sterol molecules travel through the ER, in which, given its much larger membrane surface, the sterols are diluted to a concentration below the D4H detection limit. In support of this view, BFA treatment, which strongly depletes endosomes, leads to the re-direction of sterols to other endo-membranes, including the vacuole, and likely back-flow to the PM, preventing complete D4H depletion from the PM. Thus, we favor a scenario where retrograde transport from the PM to endosomes involves passage through the ER (Fig 8).

**Figure 8.**
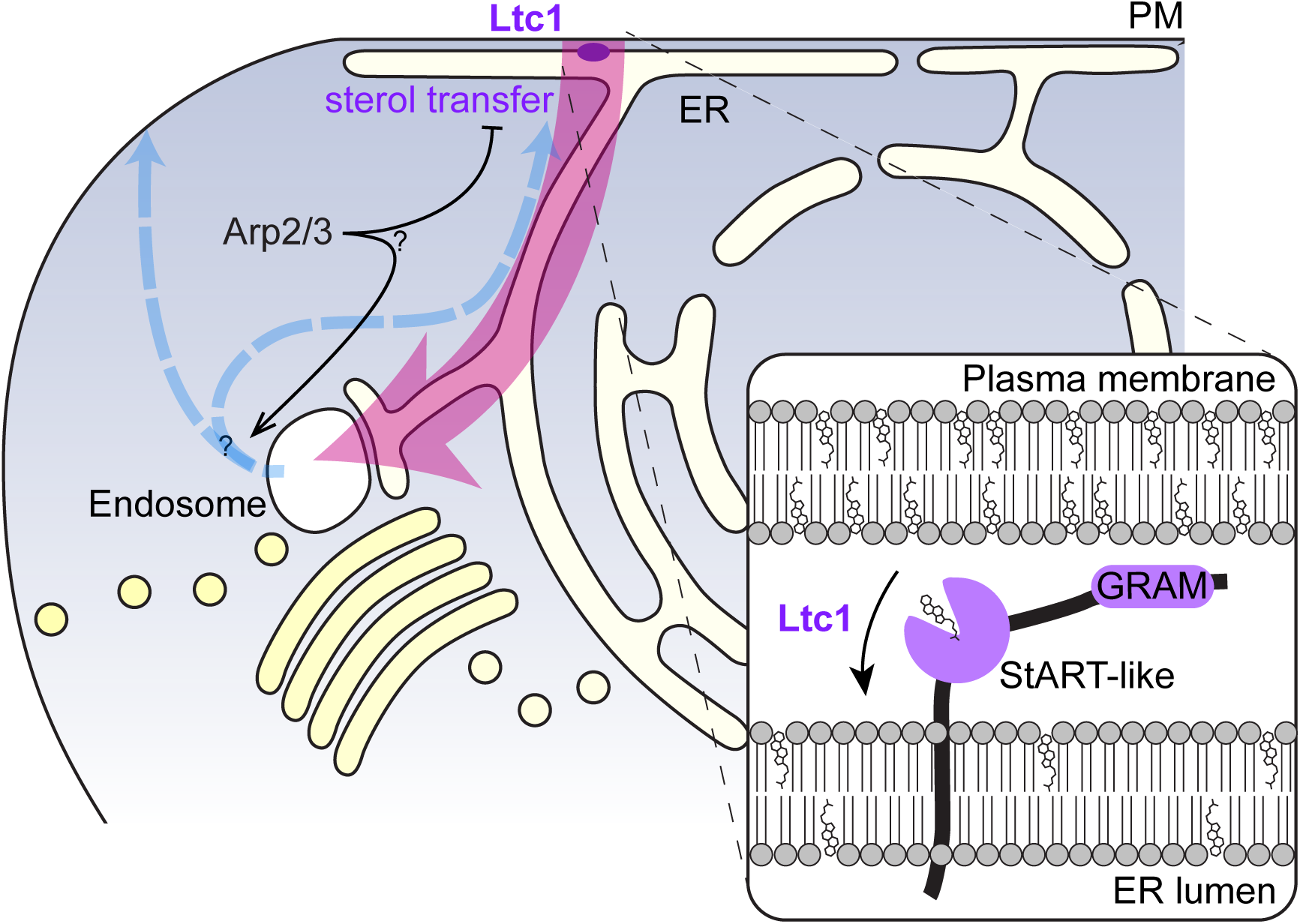
Model for the flux of sterols in fission yeast cells. Sterol molecules are transported in the cell by ways orthogonal to vesicular trafficking. In retrograde transport from the PM, the StART-like domain Ltc1 transfers sterols to the ER. Sterols then move in an anterograde manner that bypasses the Golgi to endosomes. From the endosome, sterols return to the PM independently of vesicular secretion, either through the ER or more directly. Arp2/3-dependent F-actin assembly actively promotes the transport and/or retention of sterols to the PM.

### Ltc1 is essential for retrograde sterol transport from the PM

The scenario described above implies at least two steps in PM to endosome transport: a retrograde movement between the PM and the ER, and an anterograde transport from the ER to endosomes. Three pieces of data indicate that Ltc1 mediates the first of these two steps. First, we provided several lines of evidence that the PM of *ltc1Δ* cells displays elevated sterol levels, even in unperturbed conditions. Second, Ltc1’s StART-like domain is essential for its function in retrograde sterol transport. Although we have not tested whether this domain directly binds sterols, the finding that the AmB sensitivity of *ltc1Δ* can be rescued by *S. cerevisiae* Lam5 and partly Lam6, which not only bind sterol but also shuttle it between membranes *in vitro* (Gatta et al., 2015; Murley et al., 2015), is supportive of Ltc1 directly binding and transferring sterol molecules. Third, Ltc1 localizes principally to VAP-family-independent ER-PM contact sites, at the ideal location to promote sterol retrograde flow. This localization is however not strictly required, as overexpression of cytosolic Ltc1^ΔTM^ restores sterol flow to *ltc1Δ* cells, similar to what was observed for Lam2/Ysp2 in *S. cerevisiae* (Gatta et al., 2015). This suggests that its localization at ER-PM contacts serves to increase local concentration and boost the efficiency of sterol transport. Fourth, D4H depletion from cells poles in *ltc1Δ* is consistent with the view that sterol mobilization from the ER-proximal PM at cell sides fails in this mutant, leading to sterol accumulation at cell sides. We conclude that Ltc1 mediates PM to ER retrograde sterol transfer (Fig 8).

The second step – ER to endosome transport – likely also involves direct sterol transfer at ER-endosome contacts. This is suggested from the observation that sterols could transfer to residual Syb1-compartments after BFA treatment that led to complete Golgi collapse and is consistent with previous data on sterol anterograde transport being independent of canonical vesicular trafficking system (Baumann et al., 2005; Heino et al., 2000; Urbani and Simoni, 1990). ER-endosomes contacts are well established in mammalian cells, where they function in cholesterol transfer, endosome positioning, fission and signaling (Raiborg et al., 2015). For instance, STARD3 and ORP-family ORP1L contribute to sterol trafficking from the ER to the endosome (Eden et al., 2016; Wilhelm et al., 2017). In yeast, ER-endosomes contacts have not been studied but are suggested from the Lam5 localization in *S. cerevisiae* (Weill et al., 2018). In *S. pombe*, the responsible LTP for ER-to-endosome transfer is unknown. Because, except for *ltc1Δ*, no single OSH or LAM-family deletion failed to accumulate sterols in endosomes, none of these proteins participates as sole player. The misdirection of D4H to vacuoles in *osh42Δ* mutants suggests Osh42 could promote transfer to endosomes, perhaps in conjunction to its paralog Osh41. Ltc1 could also potentially mediate sterol transfer at endosomes, as it localizes to additional locations than ER-PM contacts, and is functionally rescued by *S. cerevisiae* Lam5. In this case Ltc1 may represent an evolutionarily ancient LTP, structurally similar to, and functionally rescued by, Lam5, but whose localization and function in retrograde sterol transport from the PM are reminiscent of those of Lam2/Ysp2 (Gatta et al., 2015), for which there is no *S. pombe* orthologue. One important question for the future will be to understand what defines the directionality of the sterol transfer.

One striking consequence of blocking both endocytosis by Arp2/3 inhibition and sterol retrograde flow by *ltc1* deletion is the formation of very long PM invaginations. We hypothesize that ongoing secretion and/or transfer of other lipids in these conditions leads to PM expansion, which acquires an excessive surface accommodated in eisosomes, which expand and change shape, forming long invaginations. This observation is in line with eisosomes buffering PM size also in other stressful conditions (Kabeche et al., 2015a). Whether and how these long invaginations contribute to signaling remains to be established, but it is interesting that they appear to be in tight contact with the ER. In the presence of Ltc1, the retrograde flow of sterol, which may involve the transport of over 50 million sterol molecules at a rate of 28’000/s, similar to rate estimation in *S. cerevisiae* (Dittman and Menon, 2017), compensates for the secretion-mediated increase in membrane. Thus, retrograde sterol flow not only regulates the PM sterol concentration but also PM surface homeostasis.

### Endosome to PM transport of sterols is independent of the vesicular trafficking pathway

Like PM-to-endosome sterol transport, we found that the return of sterols to the PM is independent of the canonical vesicular trafficking pathway. Indeed, disruption of the clathrin AP-1 adaptor Apm1 and the exomer complex, which promote exit from the endosome, or the exocyst complex, which is essential for the tethering of secretory vesicles with the PM, does not prevent the return of sterols from the endosome to the PM.

These findings are consistent with earlier finding in *S. cerevisiae* that sterol anterograde trafficking occurs largely normally in temperature-sensitive mutants of the yeast secretory pathway (Baumann et al., 2005). This earlier study tracked sterol by different means, extracting PM sterols with methyl-*β*-cyclodextrin, which only extracts <0.5% of total cellular sterol. Thus, the two studies, performed in distinct organisms and with different methodologies come to the common conclusion that cellular sterol transport occurs by means independent of vesicular trafficking. One open question is the specific mechanism of sterol transfer from endosomes to the PM, which could either use a route back through the ER, or a more direct transfer from endosomes to the PM (Fig 8).

### Why does Arp2/3 inhibition lead to sterol accumulation in endosomes?

The occasional D4H-labeled endosomes in unperturbed cells indicate that the sterol flux described above also happens in steady-state growth conditions. Because acute inhibition of Arp2/3 function leads to massive accumulation of sterols in endosomes, Arp2/3 activity must be required for this flux. Remarkably, although Arp2/3 was thought to function exclusively for endocytosis in yeast, mutants with defects in endocytic uptake did not cause D4H endosomal accumulation and reacted in the same way as wildtype to Arp2/3 inhibition. These data demonstrate a novel, endocytosis-independent role for Arp2/3-mediated actin assembly in yeast cells.

One major question is where Arp2/3 acts. We consider two non-mutually exclusive possibilities. First, Arp2/3 may act at the endosome to promote anterograde sterol transfer, such that its inhibition leads to a retention of sterols in endosomes. In support of this, Arp2/3 is well-established to localize to endosomes and play numerous roles in cargo sorting and membrane fission in mammalian cells (Simonetti and Cullen, 2019). The spherical shape of STRIC post CK-666 treatment, which contrasts with the more varied shapes of Syb1-positive organelles in untreated cells, also suggests that Arp2/3-dependent actin nucleation may contribute to membrane deformation on the endosome also in yeast cells. Second, Arp2/3 may act at the PM to limit the retrograde transfer by Ltc1, such that its inhibition facilitates PM to endosome flow. For instance, inhibition may change PM properties such as membrane tension, which may induce sterol transfer by Ltc1. We note however that walled fungal cells are not believed to harbour an actin cortex similar to that present in non-walled metazoan cell (Chugh and Paluch, 2018). It will be fascinating to discover this novel function of Arp2/3 in sterol homeostasis.

## Materials and Methods

### Strains, growth conditions and drug treatment

*S. pombe* strains used in this study are listed in Supplemental Table S1. Standard genetic methods and growth conditions were used. Cells were grown in Edinburgh minimal medium supplemented with amino acids (EMM-ALU) or in rich YE5S at 25°C. Temperature sensitive (ts) *arp2-1* mutant was incubated for 6h at 36°C before analysis.

The following inhibitors were used at indicated final concentration: 200μM latrunculin A (in DMSO), 500μM CK-666 (in DMSO), 25μg/ml MBC, 0.1 μg/ml terbinafine (in ethanol), 300μM brefeldin A (BFA, in ethanol). Amphotericin B (AmB) was added at final concentration 0.2 μg/ml unless otherwise indicated. For energy depletion cells were pelleted and dissolved in EMM-ALU medium containing 20mM deoxyglucose (DG) and 10μM antimycin B (in ethanol).

Latrunculin A was ordered from Enzo Life Sciences and FM4-64 from Thermo Fisher Scientific. All other chemicals were ordered from Merck.

For recovery from CK-666 treatment, cells were incubated with CK-666 for 1h, washed twice with medium and imaged on agarose pads immediately after washing.

For filipin staining the drug was added at final concentration of 5μg/ml from DMSO stock to the cells diluted in EMM-ALU medium. Cells were imaged live within maximum 5min on glass slides.

For depletion of Sec8 from the *nmt81-sec8* strain, precultures were grown in EMM-ALU lacking thiamine and then diluted to OD 0.025 in YE5S containing thiamine, as described (Bendezu and Martin, 2011). Cells were cultivated for an additional 20h at 25°C before imaging.

For measurement of cell length and width cells were stained with Calcofluor White. 1μl of Calcofluor White stock (2mg/ml) was added to 500μl cells in EMM-ALU medium. Cells were harvested immediately, resuspended in residual medium and visualized in DAPI channel. Minimum 25 cells undergoing division were evaluated in 3 independent experiments.

For visualization of FM-64 uptake, cells were stained with 8μM FM-64 dye for 2.5 min in YE5S medium and then washed 3 times with YE5S before microscopy.

For generation of deletion mutants, wild type strains were transformed with linearized deletion plasmids (based on the pFA6a backbone) containing at least 400bp of homology to the each of the gene flanking regions. For C-terminal tagging with fluorophores, cells were transformed with fluorophore tagging plasmids carrying at least 400bp of homology to the C-terminal part of the gene and 3’UTR region.

### Microscopy

Microscopy was performed by confocal spinning disc microscopy or widefield microscopy. The majority of imaging (unless otherwise stated) was performed using spinning disc microscope - DMI4000B inverted microscope equipped with an HCX PL APO 6100/1.46 numerical aperture (NA) oil objective and PerkinElmer Confocal system (including a Yokagawa CSU22 real-time confocal scanning head and solid-state laser lines). Stacks of z-series confocal sections were acquired at 0.4μm intervals using the Volocity software. Unless otherwise indicated, images shown are single plane views.

Widefield microscopy (Fig1D) was performed on a DeltaVision platform (Applied Precision) composed of a customized Olympus IX-71 inverted microscope, an UPlan Apo 100X/1.4 NA oil objective, a 4.2Mpx PrimeBSI sCMOS camera (Photometrics) and an Insight SSI 7 color combined unit illuminator. Unless otherwise indicated, images shown are single plane views.

Filipin and Calcofluor imaging (Fig1G, 2D, 6B) was done by wide field microscopy acquired with the Leica AS AF software (Leica MicroSystems, Bannockburn, IL).

Imaging of cells, unless otherwise stated, was performed on EMM-ALU pads solidified with 2% agarose.

Imaging of *sec8* mutant and FM4-64 uptake experiments were done in glass chambers coated with soybean lectin (*Glycine max* lectin, Sigma). Shortly chambers of 96-well-plates (MGB096-1-2LG-L, Matrical Bioscience) were covered with a 100μl of filtered lectin solution (100 μg/ml in water) and kept at room temperature for 4h. Afterwards, lectin was removed, plates were washed 3 times with 200μl of water and dried. In order to attach cells to the bottom of glass chamber 100 μl of cells OD 0.2 was added to plate wells and incubated for 30 min. Afterwards, excess of cells was removed and wells were washed 2 times with medium before microscopy.

### Fluorescence Image Quantification

Quantification of PM:cytosol mCherry-D4H ratio was done on single plane images. Briefly, a segmented line was drawn along the PM and the cytoplasm area was marked using 4-pixel-wide polygon selection tool. Mean fluorescence value was calculated for each selection and corrected for background fluorescence intensity. For quantification of tip:side ratio in mCherry-D4H expressing cells cell tips or cell sides were marked using 4-pixel-wide segmented line. Mean fluorescence value was calculated for each selection and corrected for background fluorescence intensity.

For cell length and width measurements on calcofluor-stained cells, a line was drawn manually across the length and width of septated cells from the middle of one tip to the other, and the length measured using the Fiji Measure tool. Figures were prepared with Fiji (Schindelin et al., 2012), Plots of Data (Postma and Goedhart, 2019) and Adobe Illustrator.

### Correlative light-electron microscopy (CLEM) and tomography

CLEM was performed essentially as described in (Kukulski et al., 2012). Briefly, 5ml of cells grown in EMM-ALU were concentrated in 500μl and 2.5μl of a 100mM CK-666 stock in DMSO was added for 45 min to 1 h. The cells were further concentrated to a thick slurry and pipetted onto a 3-mm-wide 0.1-mm-deep specimen carrier (Wohlwend type A) closed with a flat lid (Wohlwend type B) for high-pressure freezing with a Wohlwend HPF Compact 02. The carrier sandwich was disassembled in liquid nitrogen prior to freeze substitution. High-pressure frozen samples were processed by freeze-substitution and embedding in Lowicryl HM20 using the Leica AFS 2 robot as described (Kukulski et al., 2012). 300-nm sections were cut with a diamond knife using a Leica Ultracut E microtome, collected in H20 and picked up on carbon-coated 200 mesh copper grids (AGS160; Agar Scientific). For light microscopy, the grid was inverted onto a 1x PBS drop on a microscope coverslip, which was mounted onto a microscope slide and imaged on a DeltaVision platform (Applied Precision) composed of a customized inverted microscope (IX-71; Olympus), a UPlan Apochromat 100x/1.4 NA oil objective, a 4.2Mpx PrimeBSI sCMOS camera and a colour combined unit illuminator (Insight SSI 7; Social Science Insights). Images were acquired using softWoRx v.7.0 software (Applied Precision). The grid was then recovered, rinsed in H2O and dried before post-staining with Reynolds lead citrate for 10 min. 15-nm protein A-coupled gold beads were adsorbed to the top of the section as fiducials for tomography. Transmission electron micrographs (TEM) were acquired on a FEI Tecnai 12 at 120kV using a bottom mount FEI Eagle camera (4kx4k) at 6800x or 9300x magnification for correlation with the light microscopy (LM) image. For correlation, the peripheral D4H signal was used as guide to draw the cell PM on the LM image, which was used to define the image magnification and rotation required to match it to the PM on corresponding TEM images. For tomographic reconstruction of regions of interest, tilt series were acquired at 18500x magnification over a 60° to −60° tilt range where possible at 1° increments using the Serial EM software. For tomogram reconstruction, we used the IMOD software package with gold fiducial alignment (Mastronarde and Held, 2017).

### Colony Forming Unit (CFU) Assay

CFU assays were performed in order to assess the viability of cells treated with amphotericin B. Cells were cultured to OD 0.4-0.6, pelleted and resuspended to the final OD 4 in YE5S medium containing either DMSO (1%) or CK-666 (500uM). After 1h of cultivation (25°C, shaking) amphotericin B was added to the final concentration of 5μg/ml and the samples were further incubated for another hour. Afterwards cells were collected by centrifugation, washed two times with YE5S medium and diluted to final OD 0.00025. 100μl of cells was plated on YPD plates and colonies were counted after 3 days of incubation at 30°C.

### Flow Cytometry

Flow cytometry was performed on a BD Biosciences Fortessa analyser using CellQuest software. To stain for the dead cells in the population, cells were diluted in EMM-ALU medium to final OD 0.1 and 100μl of cell suspension was mixed with 900ul EMM-ALU medium containing 1μg/ml propidium iodide. After 30sec of incubation cells were analysed by flow cytometry without gating during acquisition with 10.000 cells recorded for each sample. Data were analysed using FlowJo software.

### Spotting Assay

Sensitivity to Amphotericin B was assessed by spotting 5μl of yeast culture OD 0.25 and its 5 fold serial dilutions on plates containing EMM-ALU media with indicated AmB concentration. Growth was assessed after 72h at 30°C.

### Phylogenetic analysis

Full-length sequences of all OSBP domain-containing proteins identified in the *S. pombe* and *S. cerevisiae* genome were aligned using Muscle (https://www.ebi.ac.uk/Tools/msa/muscle/) and a neighbor-joining tree without distance correction was derived.

### Estimation of the rates of sterol flow

The following considerations were made to obtain an estimate of the number of sterol molecules internalized:

- Total PM area: 2*πr*(*l* − 2*r*) + 4*πr*^2^ ≅ 125μm^2^ for a 10μm-long, 4μm-wide *S. pombe* cell;
- Total endosome surface: 50 × 4*πr*^2^ ≅ 2.5μm^2^ for 50 endosomes of diameter 127nm;
- Estimated surface of single sterol molecule: 0.25nm^2^ (Scott, 2002)
- Minimal PM sterol concentration when labelled by D4 and D4H: 30 mol%
- Maximal PM sterol concentration when not labelled by D4H: 20 mol%
- Time of internalization: 30 min (1800 s)

If the PM contains 30 mol% sterols, this represents 1.5 × 10^8^ sterol molecules per cell. A 10 mol% drop upon Arp2/3 inhibition thus represents 5 × 10^7^ sterol molecules. 50 mol% sterol on endosome represent 5 × 10^6^ molecules, i.e. 10% of the sterols depleted from the PM. As it takes about 30 min to deplete the PM from the D4H signal, this amounts to a rate of about 28’000 molecules/s.

## Acknowledgements

We thank Dr Wanda Kukulski (LMB, Cambridge) for teaching us CLEM-tomography and for discussions, the electron microscopy facility at the University of Lausanne for help with electron microscopy sample preparation and acquisition, Prof. Gregory D. Fairn for sharing plasmids containing D4/D4H coding sequence, Dr Aleksandar Vjestica for single integration vectors and discussion, Dr Tonni Grube Andersen for discussion, Dr Quian Chen for kind gift of *end4*^*Δtld*^*pan1*^*Δapw*^ yeast strain, and lab members for careful reading of the manuscript. This work was funded by an ERC Consolidator grant (CellFusion) and a Swiss National Science foundation grant (310030B_176396) to SGM.

## Author contributions

MM performed all experiments and analysis except the CLEM, which was performed by SGM. VV helped with strain construction and imaging. MM and SGM conceived the project and wrote the manuscript. SGM acquired funding.

## Supplementary Figure Legends

**Supplementary Figure 1.**
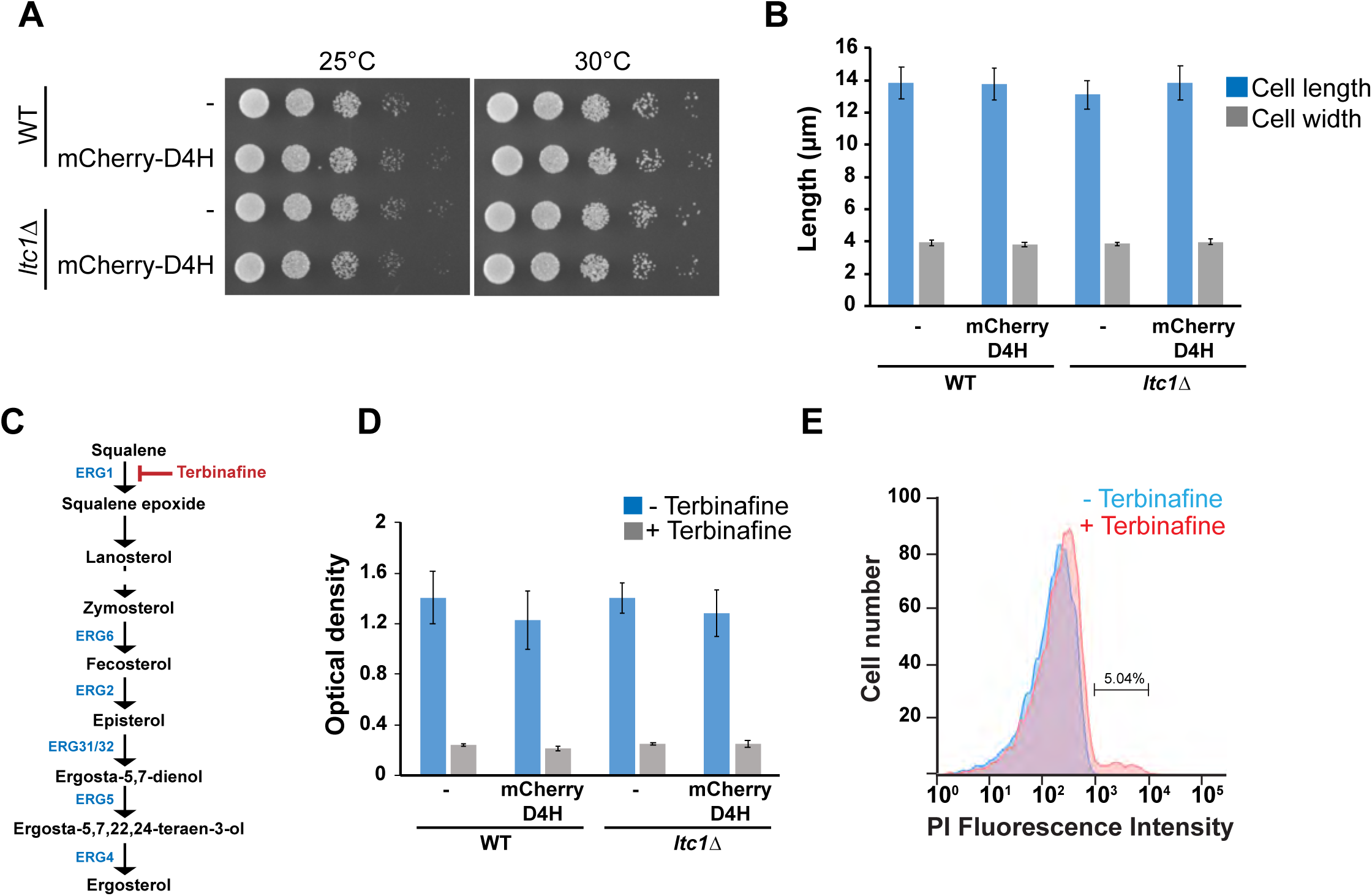
Controls for lack of D4H toxicity and terbinafine treatment. **A.** Expression of mCherry-D4H and deletion of *ltc1* do not influence growth capability of *S. pombe* cells. Serial dilutions of WT and *ltc1Δ* mutant expressing or not mCherry-D4H were compared for ability to grow at 25°C and 30°C. **B**. Expression of mCherry-D4H and deletion of *ltc1* do not influence cell length and width. WT and *ltc1Δ* mutant expressing or not mCherry-D4H were stained with calcofluor white and cell length and width of dividing cells were measured using ImageJ (n ≥ 25 cells from 3 independent experiments). **C.** Ergosterol biosynthesis pathway. **D.** Terbinafine inhibits *S. pombe* growth. WT and *ltc1Δ* mutant cells expressing or not mCherry-D4H were diluted to OD 0.02 and incubated with 0.1 μg/ml terbinafine for 25h in EMM ALU medium. Afterwards optical density was measured. Results represent three independent experiments. Error bars show the standard deviation. **E.** Terbinafine does not display cytotoxic effect in *S. pombe*. Yeast cells were treated with 0.1 μg/ml terbinafine for 16h at 25°C. Cell viability was evaluated by flow cytometry upon propidium iodide staining.

**Supplementary Figure 2.**
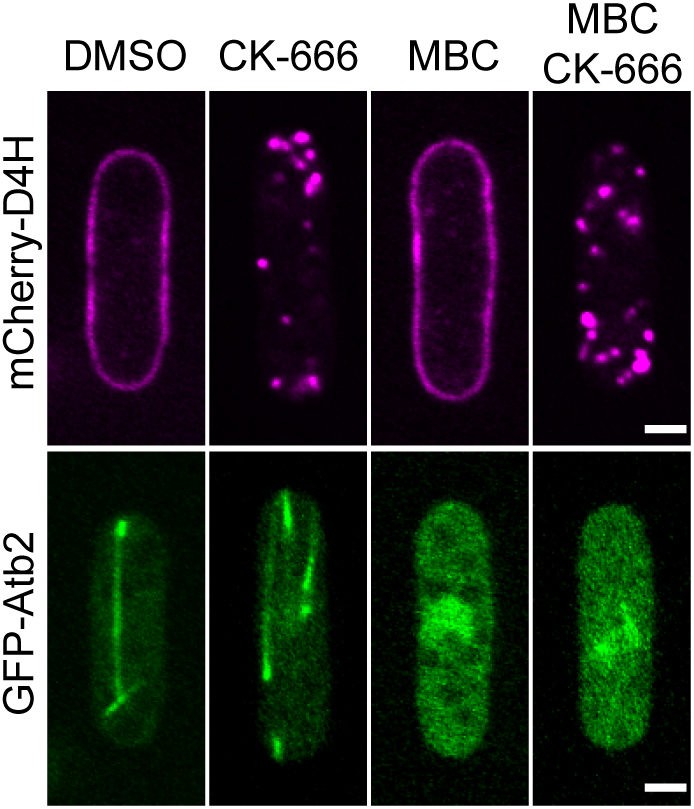
Specificity of D4H internalization and independence from microtubules. Microtubule depolymerization has no effect on mCherry-D4H distribution and relocalization. Cells expressing mCherry-D4H and tagged alpha-tubulin (GFP-Atb2) were incubated in the presence of DMSO (1%), CK-666 (500μM), methyl benzimidazol-2-yl-carbamate (MBC 25ug/ml) or were pretreated with MBC for 15min and subsequently treated with CK-666 for 1 h. Arp2/3 inhibition resulted in relocalization of mCherry-D4H signal from the PM towards the cell interior independently of microtubule integrity. Microtubule depolymerization did not trigger relocalization of mCherry-D4H. Scale bar 2μm

**Supplementary Figure 3.**
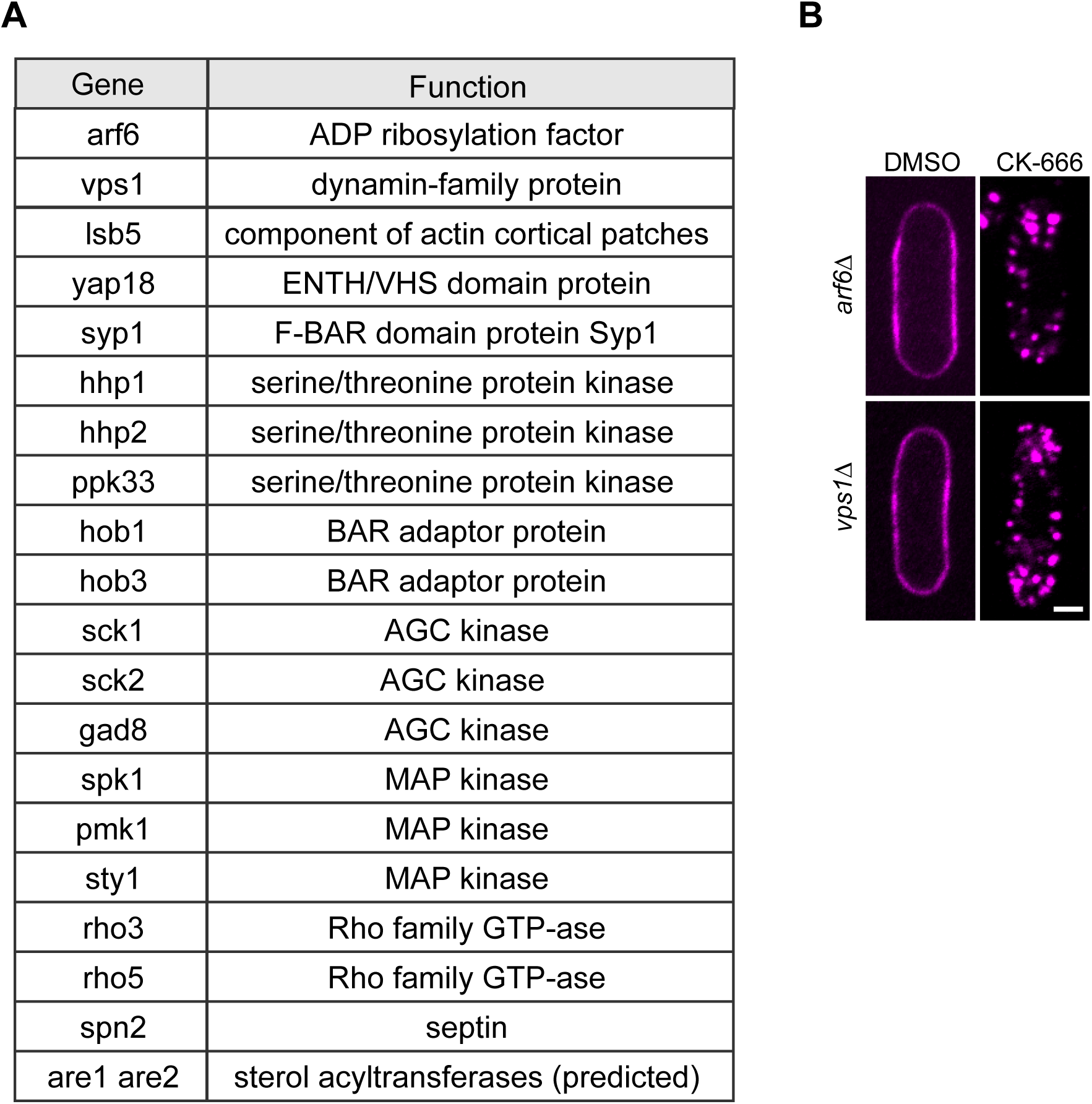
Deletion strains tested for their ability to relocate mCherry-D4H upon CK-666 treatment. **A.** Listed deletion strains expressing mCherry-D4H were treated with CK-666 for 1h and evaluated by spinning disc microscopy. All strains were capable to internalize mCherry-D4H. **B.** The deletion of *arf6* and *vps1*, encoding proteins predicted to be involved in non-clathrin mediated endocytosis, does not block mCherry-D4H internalization. *arf6Δ* and *vps1Δ* mutants expressing mCherry-D4H were incubated in the presence of 500μM CK-666 for 1h and evaluated by spinning disc microscopy. Both strains translocated sterol biosensor upon the treatment. Scale bar 2μm.

**Supplementary Figure 4.**
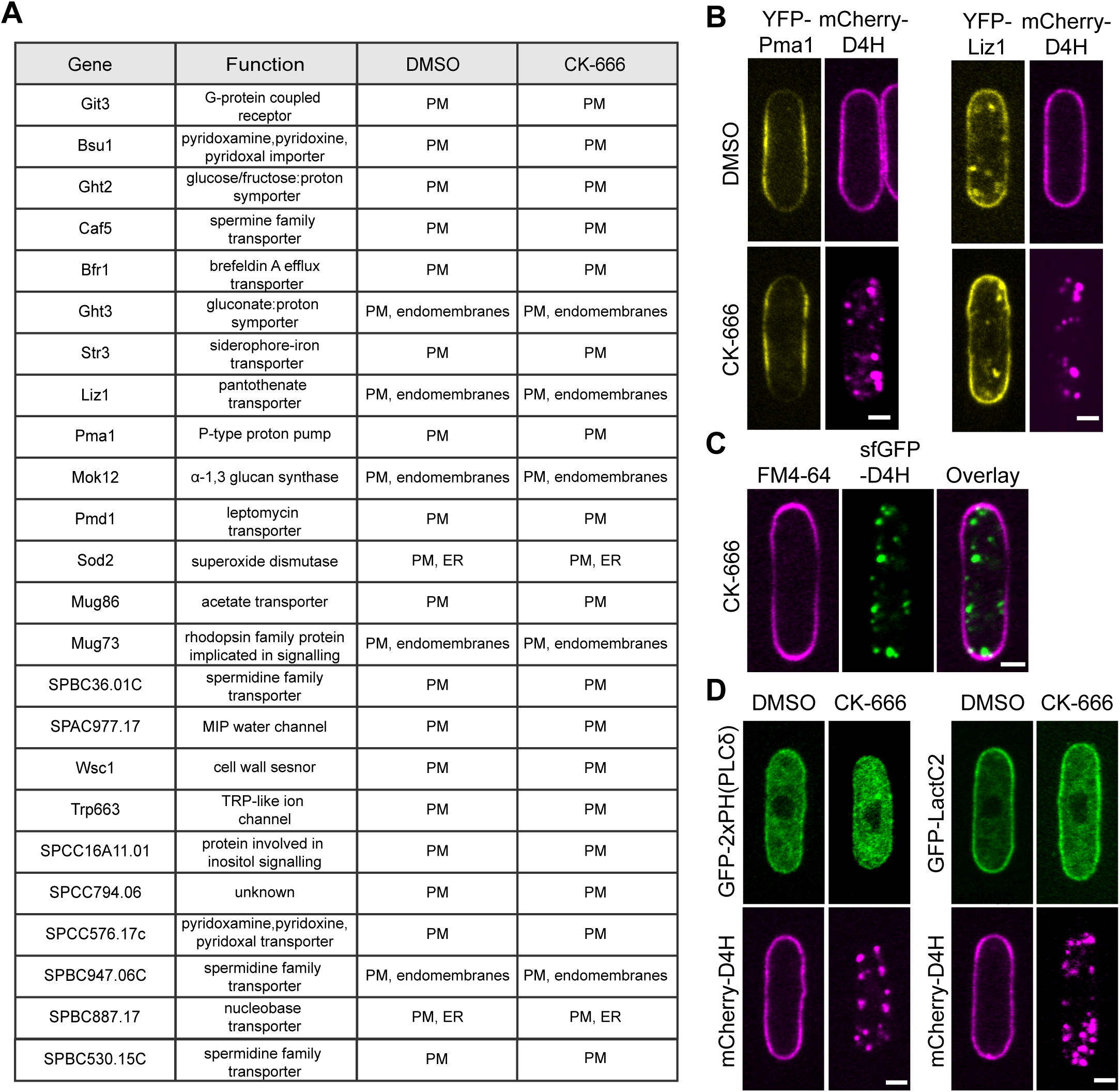
List of YFP-tagged transmembrane proteins evaluated for colocalization with mCherry-D4H positive structures. **A.** *S.pombe* cells co-expressing mCherry-D4H, which were treated with 500μM CK-666 or DMSO as control for 1h at 25°C. Localization of D4H and YFP-tagged TM proteins was evaluated by spinning disc microscopy. In all cases D4H localized to the PM in DMSO-treated cells and to internal dots in CK-666-treated ones. TM proteins could be broadly assigned to classes: those exclusively at the PM in both conditions, and those decorating both PM and internal structures (endomembranes) in both conditions. Upon strong over-expression several PM-localized TM proteins also decorated internal structures. In the second class, internal signals showed variable extent of colocalization with D4H dots upon CK-666 treatment. Notably however, none of the TM proteins was exclusively at the PM in DMSO-treated cells and in internal structures upon CK-666 treatment. We also conducted further in-depth analysis by time-lapse imaging of a couple of TM proteins of the second class (Ght3, Liz1), with the aim to capture co-internalizing signals from the PM. However, although colocalization was observed on dynamic internal structures, we did not capture a single convincing event of internalization from the PM. We conclude that the internal signal of these (and likely other) TM proteins was present prior to CK-666 treatment, and may be partly due to the natural transit of these proteins through the secretory pathway. **B.** Images of two exemplary TM proteins YFP-Pma1 localizing exclusively to PM and YFP-Liz1 localizing to PM and internal structures. Scale bar 2μm. **C.** FM4-64 does not internalize in cells treated with CK-666. **D.** Biosensors for PI(4,5)P_2_ or PS are not internalized upon CK-666 treatment. Cells expressing mCherry-D4H and either a PI(4,5)P_2_ biosensor (2xPH (PLCδ) or a PS biosensor (LactC2) were treated with 500μM CK-666 for 1h. These probes did not colocalize with D4H positive internal structures. Scale bar 2μm

**Supplementary Figure 5.**
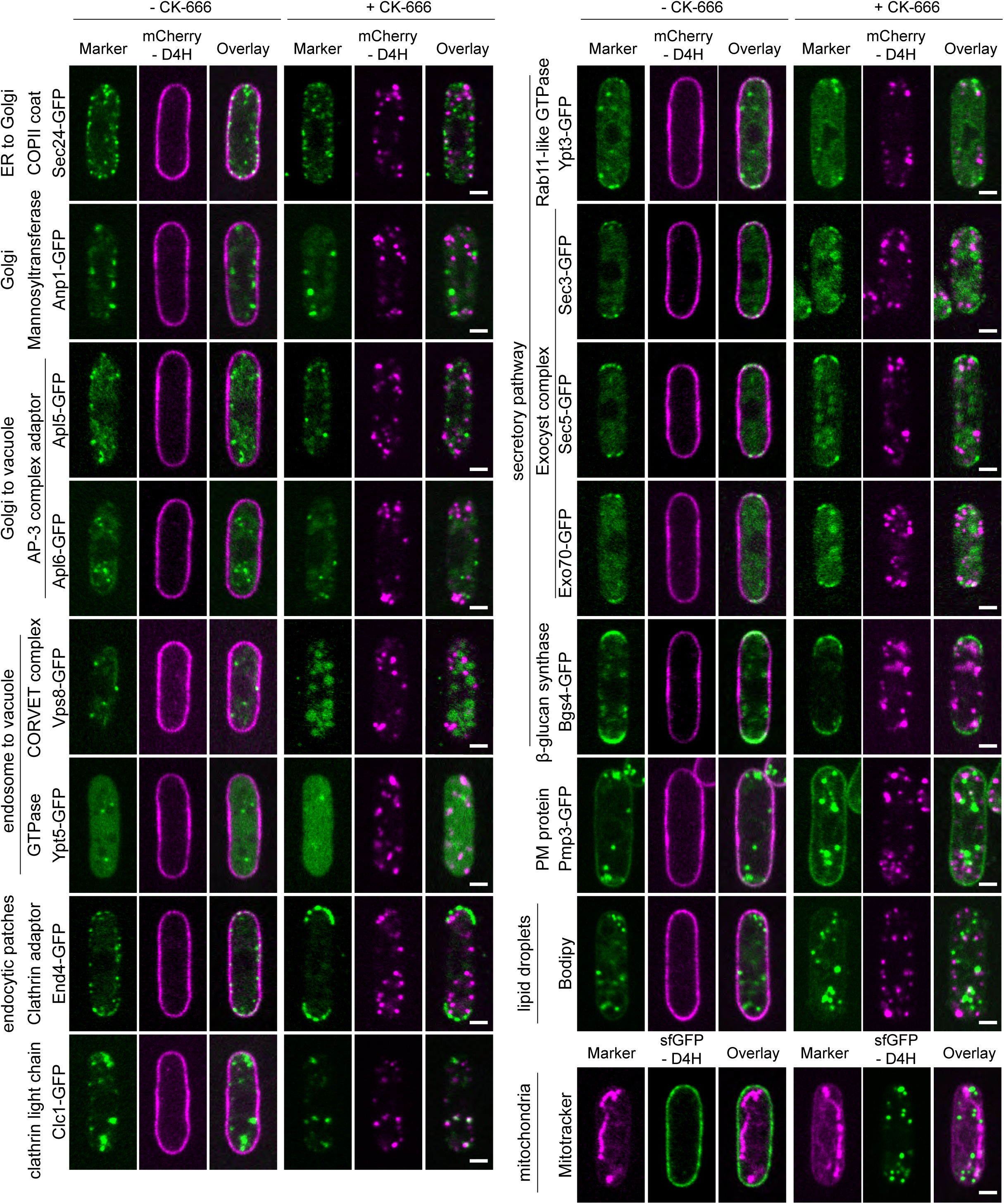
Colocalization analysis of internal D4H signal with components of endo-membranes. Marker proteins were tagged with GFP and expressed together with mCherry-D4H. Localization of both signals was compared in cells treated or not treated with CK-666. Sec24 is a COPII cargo receptor and tER marker. Anp1 is a subunit of the mannosyltransferase complex that localizes to the early Golgi. Apl5 and Apl6 are subunits of the AP3 adaptor complex present in the TGN network. Vps8 is a subunit of the CORVET complex present in the prevacuolar compartment. Ypt5 is a Rab GTPase predicted to regulate the CORVET complex. End4 is a clathrin adaptor present at sites of endocytosis. Clc1 encodes the clathrin light chain, which localizes to endosomes/TGN and actin patches. Ypt3 is a Rab GTPase homologous to Rab11, localizing to secretory vesicles and the TGN. Sec3, Sec5 and Exo70 are exocyst components, localizing to secretory vesicles and the PM. Bgs4 is a 1,3-beta-glucan synthase present at cell tips and within TGN. Pmp3 is a predicted PM protein, which also labels prominent, unknown internal structures. Bodipy labels lipid droplets. Mitotracker labels mitochondria. No colocalization was observed between any of these markers and D4H internal structures upon CK-666 treatment, except for Clc1, for which partial colocalization was observed. Scale bar 2μm

**Supplementary Figure 6.**
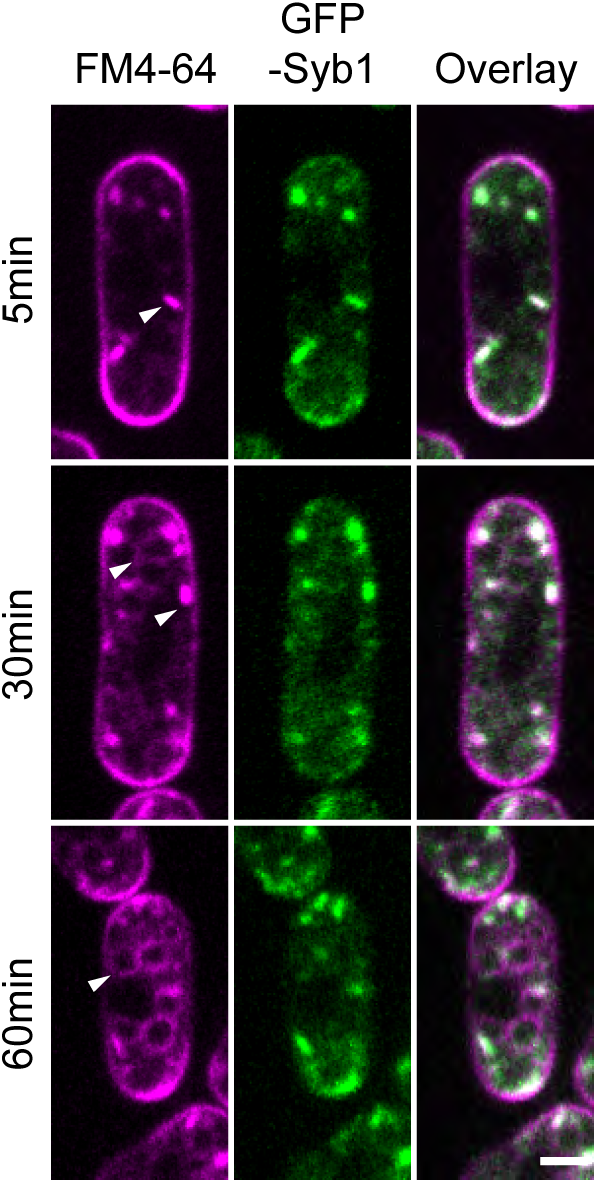
Colocalization of GFP-Syb1 with internalized FM4-64. GFP-Syb1 labels an endosomal compartment. Cells were incubated with FM-64 and dye distribution was evaluated after 5, 30 and 60min post labelling. FM4-64 signal initially labels PM and endosomes, which strongly colocalize with GFP-Syb1. Endosomal and weak vacuolar staining can be observed 30min post staining and the dye reaches vacuoles within 60min of incubation. Scale bar 2μm.

## Supplementary Movies legends

**Movie S1. mCherry-D4H distribution during vegetative growth.** Spinning-disk confocal time-lapse of mCherry-D4H during the interphase growth in *S. pombe*. The movie is sped up 1200-fold. Timing starts from the time the cells are placed on the agarose pad.

**Movie S2. mCherry-D4H relocalization upon CK-666 treatment.** Spinning-disk confocal time-lapse of mCherry-D4H after treatment with 500μM CK-666. The movie starts 4 min post CK-666 addition and is sped up 1200-fold.

**Movie S3. Recovery of mCherry-D4H cortical localization upon removal CK-666.** Spinning-disk confocal time-lapse of mCherry-D4H distribution recovery after wash-out of CK-666. The movie starts 6 min post wash-out and is sped up 600-fold.

**Movie S4. Colocalization of mCherry-D4H with Sec72-GFP during treatment with CK-666.** Spinning-disk confocal time-lapse of cells co-expressing mCherry-D4H and Sec72-GFP after treatment with CK-666. The movie is sped up 10-fold.

**Movie S5. Colocalization of mCherry-D4H and GFP-Syb1 during treatment with CK-666.** Spinning-disk confocal time-lapse of mCherry-D4H and GFP-Syb1 distribution after treatment with CK-666. The movie is sped up 10-fold.

**Movie S6. mCherry-D4H distribution during vegetative growth in *ltc1Δ* mutant.** Spinning-disk confocal time-lapse of mCherry-D4H during the interphase growth in *ltc1Δ*. The movie is sped up 1200-fold. Timing starts from the time the cells are placed on the agarose pad.

**Movie S7. mCherry-D4H relocalization upon CK-666 treatment in *ltc1Δ* mutant.** Spinning-disk confocal time-lapse of mCherry-D4H distribution upon treatment with CK-666 in *ltc1Δ*. The movie starts 4 min post CK-666 addition and is sped up 1200-fold.

**Supplementary Table S1.**
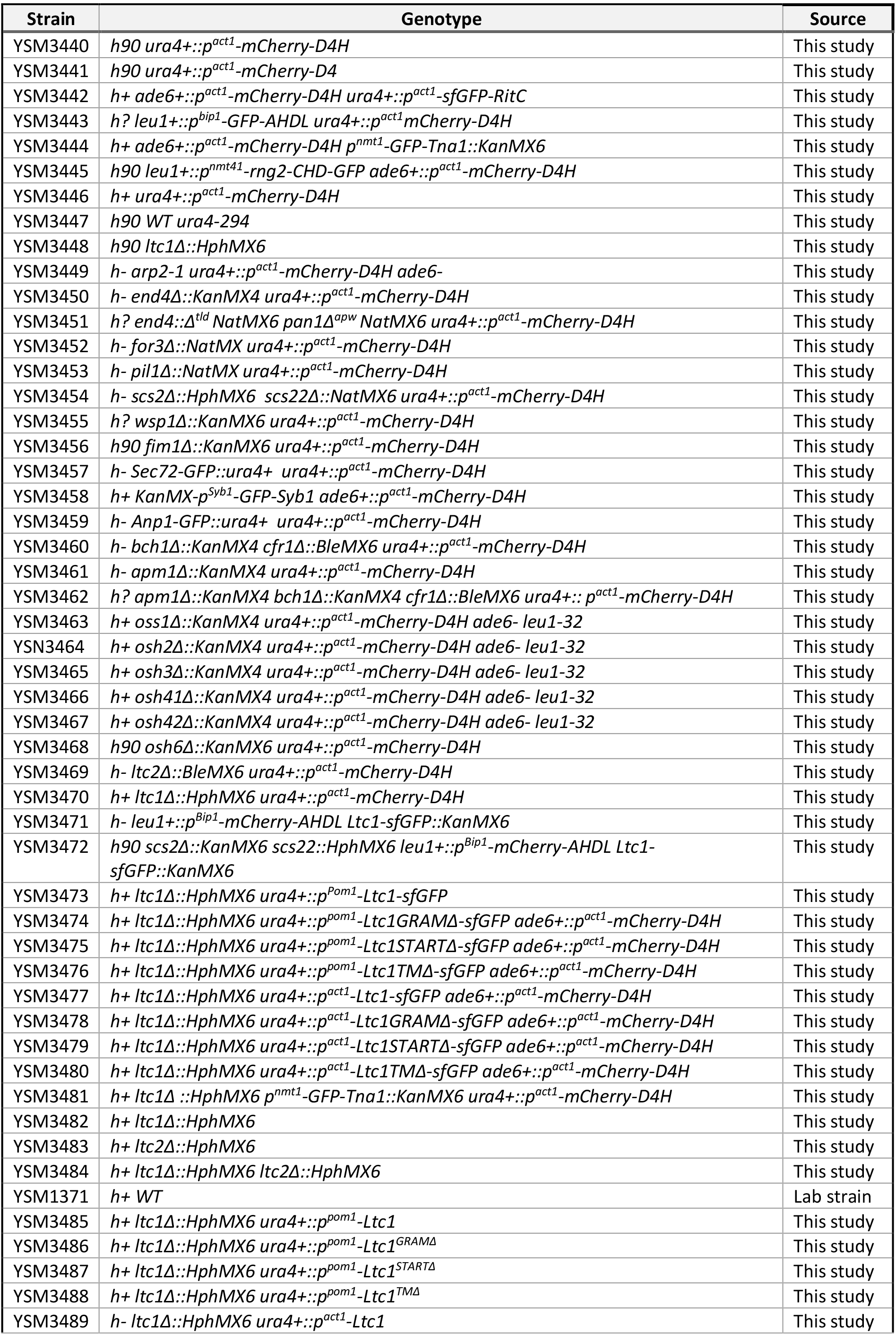
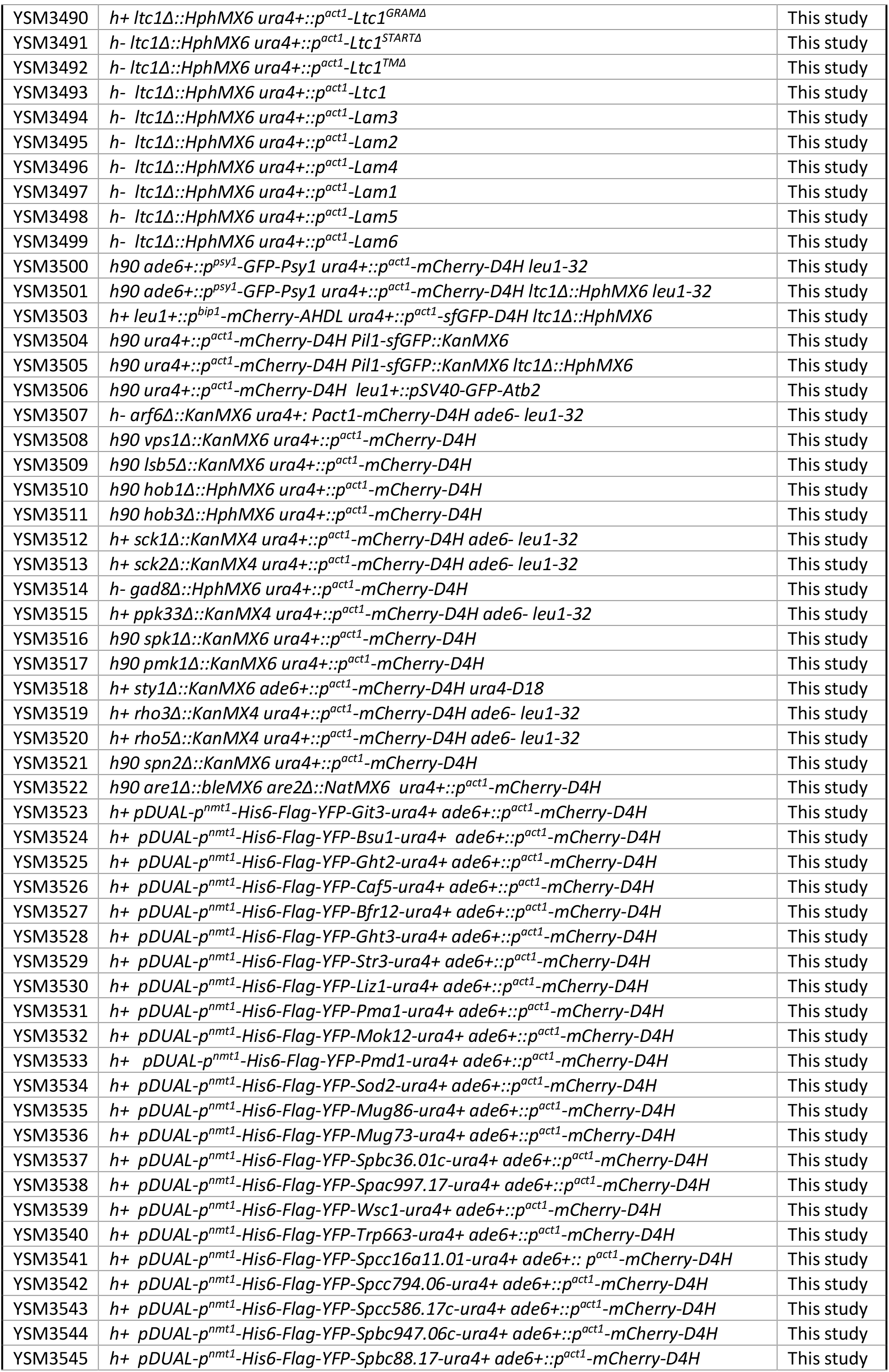
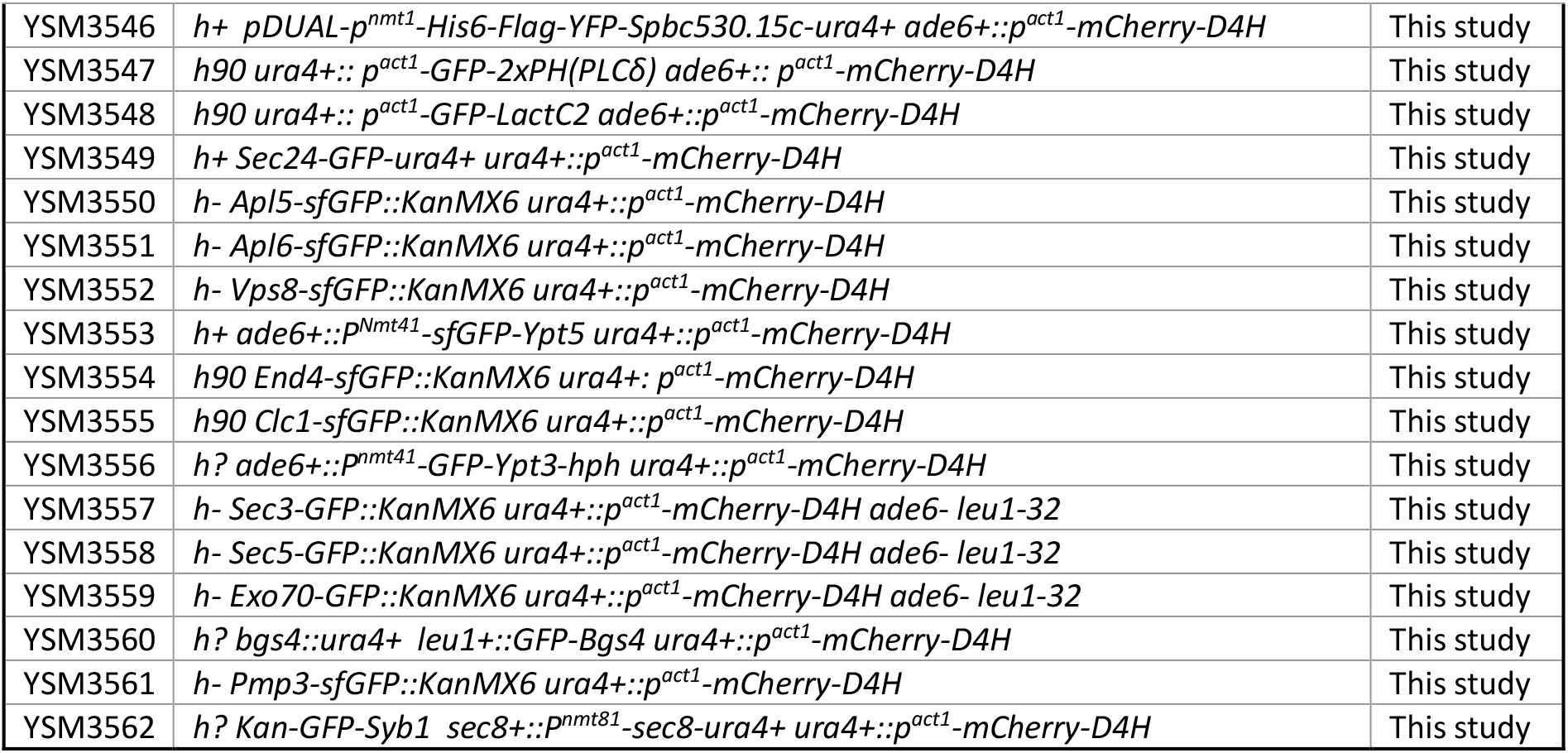

